# MitoDelta: identifying mitochondrial DNA deletions at cell-type resolution from single-cell RNA sequencing data

**DOI:** 10.1101/2025.06.10.658806

**Authors:** Haruko Nakagawa, Yasuyuki Shima, Yohei Sasagawa, Itoshi Nikaido

## Abstract

**Background:** Deletion variants in mitochondrial DNA (mtDNA) are associated with various diseases, such as mitochondrial disorders and neurodegenerative diseases. Traditionally, mtDNA deletions have been studied using bulk DNA sequencing, but bulk methods average signals across cells, thereby masking the cell-type-specific mutational landscapes. Resolving mtDNA deletions at single-cell resolution is beneficial for understanding how these mutations affect distinct cell populations. To date, no specialized method exists for detecting cell-type-specific mtDNA deletions from single-cell RNA sequencing data. Notably, mtDNA possesses unique molecular features: a high copy number, stable transcription, and compact structure of the mitochondrial genome. This results in a relatively high abundance of mtDNA-derived reads even in single-cell RNA sequencing data, suggesting the possibility of detecting mtDNA deletion variants directly from transcriptomic data.

**Results:** Here, we present MitoDelta, a computational pipeline that enables the detection of mtDNA deletions at cell-type resolution solely from single-cell RNA sequencing data. MitoDelta combines a sensitive alignment strategy with robust statistical filtering based on a beta-binomial model, allowing accurate identification of deletion events even from noisy single-cell transcriptomes. To capture cell-type-specific deletion patterns, MitoDelta analyzes reads pooled by annotated cell types, enabling quantification of deletion burden across distinct cellular populations. We benchmarked MitoDelta against existing mtDNA deletion detection tools and demonstrated superior overall performance. As a practical application, we applied MitoDelta to a published single-nucleus RNA sequencing dataset for Parkinson’s disease and revealed distinct mtDNA deletion burdens across neuronal subtypes.

**Conclusions:** MitoDelta enables the transcriptome-integrated, cell-type-specific detection of mtDNA deletions from single-cell RNA sequencing data alone, offering a valuable framework for reanalyzing public datasets and studying mitochondrial genome alterations at cell-type resolution. This integrated approach enables insights into how mtDNA deletions are distributed across specific cell types and cellular states, providing new opportunities to investigate the role of mtDNA deletions in cell-type-specific disease mechanisms. The tool is available at https://github.com/NikaidoLaboratory/mitodelta.

## Background

Mitochondria possess their own genome, which is highly prone to mutations. Variants in mitochondrial DNA (mtDNA) are associated with various diseases, including mitochondrial diseases, neurodegenerative disorders, and cancers [1–4]. In particular, large-scale mtDNA deletions have been reported in neurodegenerative diseases such as Parkinson’s disease and Alzheimer’s disease [5], as well as in congenital disorders such as Kearns–Sayre syndrome, Pearson syndrome, and chronic progressive external ophthalmoplegia [6]. While some mtDNA variants are maternally inherited, others arise somatically. For example, somatic mutations are implicated in impaired DNA error-correction mechanisms, such as function-changing mutations in mtDNA polymerase genes like *POLG* [7,8]. Accurate calling of mtDNA deletions is therefore helpful for understanding their roles in disease pathologies.

Traditionally, mtDNA deletions have been studied by bulk DNA sequencing (DNA-seq), such as targeted deep sequencing or whole-genome sequencing (WGS) [9–11]. These methods are robust and reliable but are not well suited for investigating disease or cellular heterogeneity. In other words, bulk methods average signals across cells, masking cell-type-specific mutational landscapes. Researchers have employed cell isolation techniques (e.g., cell sorting with flow cytometry and laser-capture microdissection) to prepare specific cell types before sequencing to address this issue. However, these low-throughput approaches are labor intensive and typically do not provide accompanying transcriptomic information.

Recent advances in single-cell technologies now allow high-throughput mtDNA variant analysis at single-cell resolution. Single-cell DNA sequencing (e.g., single-cell WGS and single-cell whole-exome sequencing) can be used for mtDNA variant calling at the single-cell level [12]. While these approaches are the most straightforward methods to detect variants at single-cell resolution, their high cost limits their widespread use.

Additionally, targeting the whole genome is not economical when researchers are interested in only the mitochondrial region. Alternative mtDNA-specific single-cell approaches have been developed to solve this issue [13], including MitoSV-seq [14], MAESTER [15], mtscATAC-seq [16,17], DOGMA-seq [18], and ReDeeM [19]. These methods are effective for mtDNA-focused analyses but require specialized protocols, such as mtDNA enrichment or deep sequencing. Additionally, except for mtscATAC-seq, these methods target only the genome, not the transcriptome, making it challenging to know the functional state of the corresponding cells. These current methods are less suitable for a variant analysis in a transcriptomic context.

On the other hand, several tools have been developed to infer mtDNA variants from single-cell RNA sequencing (scRNA-seq) data [20,21]. They do not require additional experiments, providing insights into both the mutations and their transcriptional consequences. However, most of them are designed for single-nucleotide variants (SNVs) or indels, not large deletions. For mtDNA deletion variant calling, tools such as MitoSAlt [22], eKLIPse [23], and Splice-Break [24,25] are available, but these tools are designed primarily for mtDNA-enriched DNA-seq or WGS data, and their applicability to scRNA-seq remains limited. To date, no tools currently exist to detect cell-type-specific mtDNA deletions solely from scRNA-seq data.

Although often overlooked, detecting mtDNA deletion variants from scRNA-seq data is an attractive approach for two reasons. First, mtDNA has unique molecular characteristics. It is a circular molecule that is present in high copy numbers and encodes mRNA, tRNA, and rRNA with minimal intergenic regions and no introns. This structure results in an almost continuous read distribution across the mitochondrial genome, with very few gaps [26]. As a result, even scRNA-seq data, which generally have limited coverage, may contain sufficient reads for variant calling in the mitochondrial region. Second, scRNA-seq enables simultaneous access to mutational and transcriptomic information, providing insights into the potential phenotypic effects of mtDNA deletions. This analysis enables us to explore not only the mutation landscape across different cell types but also the potential impact of mtDNA deletions on the cellular state. However, the use of scRNA-seq data for detecting mtDNA deletions presents two major challenges. First, the limited number of reads per cell makes sensitive variant detection difficult, increasing the risk of false negatives. Second, limited coverage also increases the likelihood of false positives, making reliable detection challenging.

We developed MitoDelta (Mitochondrial genome DELetions in cell-Type Analysis), a computational pipeline for robust and reliable cell-type-specific mtDNA deletion variant calling, to address these challenges. MitoDelta employs LAST, a fast and sensitive local aligner [27], to ensure high recall performance. Furthermore, we incorporated statistical filtering via a beta-binomial distribution model to suppress false positives. It reduces the number of technical artifacts and improves the specificity and confidence in variant calling. In this study, we systematically benchmarked MitoDelta against three published deletion callers using both simulated and real data, demonstrating MitoDelta’s superior performance in analyzing scRNA-seq data. We also evaluated its effectiveness by analyzing a public dataset of Parkinson’s disease patients [28], verifying its ability to reveal the deletion landscape at cell-type resolution. MitoDelta provides insights into the integrity of the mitochondrial genome and its role in disease.

## Methods

### MitoDelta: accurate mtDNA deletion variant identification from scRNA-seq data at cell-type resolution

MitoDelta is a computational pipeline for calling mtDNA deletion variants at cell-type resolution from scRNA-seq data. The pipeline consists of three main steps: (1) read pooling by cell type, (2) candidate deletion variant calling, and (3) statistical filtering of variants.

### Data preprocessing

Before running MitoDelta, users should preprocess scRNA-seq data using standard workflows (e.g., CellRanger) to generate two files: a BAM file containing aligned reads and a cell-by-gene expression matrix. MitoDelta is compatible with data from 10x Genomics Chromium and other scRNA-seq platforms, as long as they use cell barcode (CB)-based labeling. The cell type annotation should be performed by analytical tools (e.g., Seurat, Scanpy), and the resulting cell type assignments should be exported in tab-separated value (TSV) format, in which each row contains a CB and its corresponding cell type label. This file serves as input for the cell-type-specific analysis in MitoDelta.

### Read pooling and variant calling

In the initial step of MitoDelta, the original multiplexed BAM file is split into cell-type-specific BAM files based on the cell annotations using a custom Python script [29]. These cell-type-split BAM files are then converted to FASTQ format by SAMtools (v1.12). In the second step, the resulting FASTQ files are processed by a variant calling process based on LAST (v2.34), using a strategy inspired by MitoSAlt [22].

### Statistical filtering

In the final step, MitoDelta applies error-model-based statistical filtering to reduce false-positive calls, which are particularly prevalent in shallow scRNA-seq data. MitoDelta models background sequencing noise using a beta-binomial distribution, which accounts for the overdispersion typically observed in low-coverage sequencing datasets [30,31]. The model is defined as:

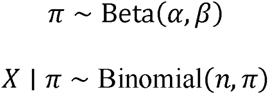

Here, π represents the true error rate, and *X* is the observed number of breakpoint-supporting reads among *n* total reads. The parameters α and β are estimated using the method of moments, based on background noise-derived calls. Specifically, variants with heteroplasmy levels below 1% are empirically assumed to be artifacts and are used to fit the model. We use the average (μ) and variance (σ^2^) of the background signal to estimate the parameters as follows:

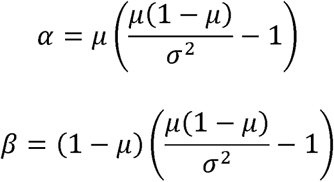

Once the beta-binomial model is fitted, each candidate variant is evaluated by calculating the probability of its observed support under the null hypothesis that it arises from background noise. Before this calculation, we excluded variants supported by only a single breakpoint read or lacking wild-type reads to minimize the influence of ultralow-read-depth regions. P values are then computed for the remaining candidates and adjusted for multiple testing using the BenjaminiLHochberg procedure. Variants with an adjusted p value (q value) less than 0.05 are considered statistically significant and retained as high-confidence mtDNA deletion variants.

### Quantitative benchmarking of MitoDelta and existing mtDNA deletion callers with simulated RNA-seq data

We evaluated the performance and computational time of mtDNA deletion detection on a simulated RNA-seq dataset for MitoDelta and three existing mtDNA deletion callers: MitoSAlt (v1.1.1) [22], eKLIPse (v1.8) [23], and Splice-Break2 (v3.0.1) [24,25]. All tools were tested using their default settings, with the exception that the minimum required number of breakpoint-supporting reads was set to one across all tools to ensure fair comparisons.

Simulated reads were generated using Flux-simulator (v1.1) [32], an *in silico* tool that produces RNA-seq reads based on a reference genome and corresponding transcript information. We used the mouse mitochondrial genome (NC_005089.1) and its RefSeq-based transcript annotations (mm39) in FASTA and GTF formats. The positions of these genes were assigned semirandomly across the mitochondrial genome to assess the performance on various deletions. We predefined ten deletion lengths (50, 120, 250, 500, 1,000, 2,000, 4,000, 6,000, 8,000, and 10,000 bp), and for each length, ten genomic coordinates within the mitochondrial genome were randomly selected to generate 100 unique deletion events. Using custom Python scripts, we created corresponding modified FASTA and GTF files, each representing one deletion variant. Then, RNA-seq reads were simulated for each modified genome using the following Flux-simulator parameters: POLYA_SHAPE = 2, POLYA_SCALE = 80, RTRANSCRIPTION = YES, RT_PRIMER = PDT, RT_LOSSLESS = YES, RT_MIN = 150, RT_MAX = 7500, FRAG_METHOD = NB, FRAG_SUBSTRATE = DNA, FILTERING = YES, READ_LENGTH = 100, PAIRED_END = NO, READ_NUMBER = 50000, FASTA = true, and PCR_DISTRIBUTION = none. From the 100 simulated deletion events, we selected 20 variants with relatively high read support as the “true” deletions. A total of 50,000 reads derived from the deletion-containing reference were mixed with 200,000 reads from the wild-type reference to simulate heteroplasmy. The remaining 80 deletion events, which were supported by very few reads due to low coverage in the simulation, were used as background noise. Reads corresponding to these 80 deletions were merged and downsampled to 250,000 reads to emulate sequencing noise, which is commonly observed in real data. These error-like reads were then added as noise to each of the evaluation datasets, resulting in 500,000 reads per deletion. Finally, each dataset was downsampled to five different coverage levels (5,000, 10,000, 50,000, 100,000, and 500,000 reads) using seqkit (v2.9.0). This range was chosen based on two assumptions. First, one of the most common scRNA-seq platforms, 10x Genomics Chromium, recommends coverage of approximately 20,000 reads per cell [33]. Second, although the proportion of mitochondrial reads varies across tissues, we conservatively assumed it to be approximately 5%, meaning that each cell would contribute approximately 1,000 mitochondrial reads. We simulated datasets representing pooled inputs from 5 (5,000 reads), 10 (10,000 reads), 50 (100,000 reads), 100 (200,000 reads), and 500 cells (500,000 reads) to evaluate our cell pooling approach, with particular emphasis on the 50-cell pool and its surrounding conditions. In total, 20 distinct deletion variants across five coverage levels were simulated, and the calling performance was evaluated in terms of the recall, precision, accuracy, and F1 score.

### Comprehensive benchmarking of MitoDelta and existing mtDNA deletion callers on simulated and real scRNA-seq data

#### Customized workflows of existing tools by recombining components

For a more comprehensive evaluation of mtDNA deletion callers, we benchmarked MitoDelta and three existing tools on both simulated and real RNA-seq datasets. In addition to comparing the default workflows of these tools, we systematically tested customized workflows by recombining components at each step. MitoSAlt and eKLIPse share a similar three-step structure: premapping, read selection for the alignment step, and local alignment for deletion detection. For these tools, we tested combinations of three mapping tools for premapping (BWA-MEM2 (v2.2.1), HISAT2 (v2.2.1), or STAR (v2.11b)), two types of reference genomes used for premapping (mtDNA only or the whole genome), and two local alignment tools (BLAST (v.2.2.28) or LAST (v2.34)), resulting in 12 different method combinations. In the premapping step, we used the reference genome downloaded from UCSC (mm39 for mouse and hg38 for human). For mouse data, we observed that nuclear mitochondrial DNA (NUMT), segments of DNA sequences transferred from mtDNA into the nuclear genome, can erroneously attract mtDNA-derived reads. We used an NUMT-masked reference genome for the mouse data to minimize these false alignments. Splice-Break2 employs MapSplice to identify splice junction candidates, which are then used as breakpoint-supporting reads. Thus, we tested two mapping strategies, MapSplice and STAR, yielding two workflow variants for this tool. All customized workflows, together with MitoDelta, were benchmarked on both simulated and real scRNA-seq datasets to assess their qualitative and quantitative performance.

#### Simulated RNA-seq data

For benchmarking, we used the same simulated RNA-seq data described earlier. This dataset included synthetic reads generated from a reference mtDNA sequence containing 20 predefined deletions. The true heteroplasmy levels of each deletion were manually curated using the Integrative Genomics Viewer (IGV) [34] (v2.19.4) to evaluate the accuracy of the heteroplasmy estimate. These curated values served as the ground truth for the quantitative assessment.

#### Real scRNA-seq data from mtDNA deletion-associated disease samples

As a real dataset for benchmarking, a published scRNA-seq dataset from peripheral blood mononuclear cells (PBMCs) of Pearson syndrome patients (GSE173932) was used [35]. The dataset included samples from three patients, each harboring a distinct mtDNA deletion: Patient 1 (deletion: 6072–13095, heteroplasmy: 41%), Patient 2 (deletion: 8469–13446, heteroplasmy: 66%), and Patient 3 (deletion: 10381–15406, heteroplasmy: 54%). These reference heteroplasmy values were obtained from the original study. Additionally, RNA-level heteroplasmy was manually investigated using IGV. We prepared the benchmark dataset in a pseudobulk manner by downsampling the original FASTQ files to seven coverage levels to assess the effect of the sequencing depth: 20,000, 100,000, 200,000, 1,000,000, 2,000,000, 10,000,00, and 20,000,000 reads. This range was determined based on the recommended sequencing depth by the Chromium platform, simulating the input from a single cell up to pools of 1,000 cells. Among these, the coverage level corresponding to a 50-cell pool (∼1,000,000 reads) was a primary target, and surrounding depths were included to evaluate its robustness. Each coverage level was replicated five times with different random seeds.

### Analysis of cell-type-specific mtDNA deletion variants in data from patients with Parkinson’s disease via MitoDelta

#### Data preprocessing

The single-nucleus RNA-seq (snRNA-seq) data from the postmortem brain tissues of Parkinson’s disease (PD) patients and control individuals were obtained from GSE243639 [28]. Twenty-nine samples from the substantia nigra pars compacta of 14 controls and 15 PD patients were sequenced using the 10x Genomics Chromium platform. Each patient’s sample was registered across multiple runs in the database, and thus the FASTQ files were merged by patient using the cat command. Merged FASTQ files were then preprocessed using CellRanger (v8.0.1) with the GRCh38-2020-A-2.0.0 reference genome [36].

#### Quality control, data integration, clustering, and cell type annotation

The single-cell analysis of the cell-by-gene expression matrix was performed with the Seurat (v5.0.0) R package [37]. The datasets were converted into Seurat objects with the Read10x() function. Quality control was conducted using the method described in the original study, excluding nuclei with a low RNA content (< 500 transcripts) or high RNA content (> 10,000 transcripts). Additionally, MALAT1 was removed from the transcript list because of its high enrichment, which could introduce bias into clustering. The mitochondrial RNA content was filtered at a threshold of 15%. Normalization and principal component analysis (PCA) were applied to each dataset using the SCTransform() (vst.flavor = v2) and RunPCA(), respectively. Integration features were selected using the SelectIntegrationFeatures() function with 6,000 features.

FindIntegrationAnchors() identified integration anchors, with sample s.0096 as the reference, employing the canonical correlation analysis (CCA). The datasets were subsequently integrated using the IntegrateData(). Following integration, PCA was repeated on the integrated dataset, and the top 20 principal components were used for dimensionality reduction via UMAP using the RunUMAP() function. A shared nearest neighbor graph was constructed with the FindNeighbors() function on the first 20 PCs, and clustering was performed with the FindClusters() function at a resolution of 0.6.

Clusters were annotated based on marker gene expression, classifying cells into major cell types, including astrocytes, microglia, neurons, oligodendrocytes, oligodendrocyte precursor cells (OPCs), T cells, and vascular cells (VCs). Neurons were further classified into dopaminergic, GABAergic, and glutamatergic subtypes based on their transcriptomic profiles. Clusters with extremely low cell numbers, sample-specific biases, or a lack of distinctive gene expression patterns were excluded. CBs and corresponding cell type annotations were then saved in TSV format.

#### Data splitting, variant calling, and filtering in the MitoDelta workflow

Multiplexed BAM files were split by cell type based on the provided cell-type annotations. Up to 50 cells per cell type per sample were randomly subsampled to ensure a fair evaluation across samples and cell types, and the cell-type-specific BAM files were converted to FASTQ format by SAMtools. Then, the resulting cell-type-specific FASTQ files were processed through MitoDelta.

The obtained deletion calls were evaluated from two perspectives. First, we examined whether MitoDelta detected the common mtDNA deletion reported in PD patients (chrM: 8469-13447). These calls were also visually confirmed using IGV. Second, we compared cumulative deletion burdens across cell types by performing the KruskalLWallis test for PD patients and control individuals. When significant differences were observed, we subsequently performed Dunn’s test, with correction for multiple testing via the BenjaminiLHochberg procedure.

## Results

### The workflow of MitoDelta

MitoDelta is a computational pipeline designed to identify mtDNA deletion variants at cell-type resolution from scRNA-seq data (Fig. 1A). The workflow consists of three key steps: (1) read pooling by cell type, (2) detection of candidate breakpoints via local alignment, and (3) statistical filtering. In the first step, reads are split and pooled by cell type. This approach serves two purposes: enabling a cell-type-resolved analysis and ensuring sufficient read coverage, which is often too sparse at the individual cell level. In the second step of detecting breakpoint-supporting reads, MitoDelta employs LAST, a sensitive local aligner. Although its computational cost is higher than that of standard mapping tools, we consider it feasible for shallow scRNA-seq data with reasonable computational efficiency. However, its sensitivity comes with a trade-off of many false positives. The third step incorporates a beta-binomial distribution model that is used for modeling the overdispersion of scRNA-seq or low-abundance DNA-seq data to solve this issue. The model allows MitoDelta to robustly distinguish true deletions from background noise. By combining sensitive breakpoint detection with statistical filtering, we aimed to achieve balanced recall and precision, even in sparse scRNA-seq data.

**Figure 1.**
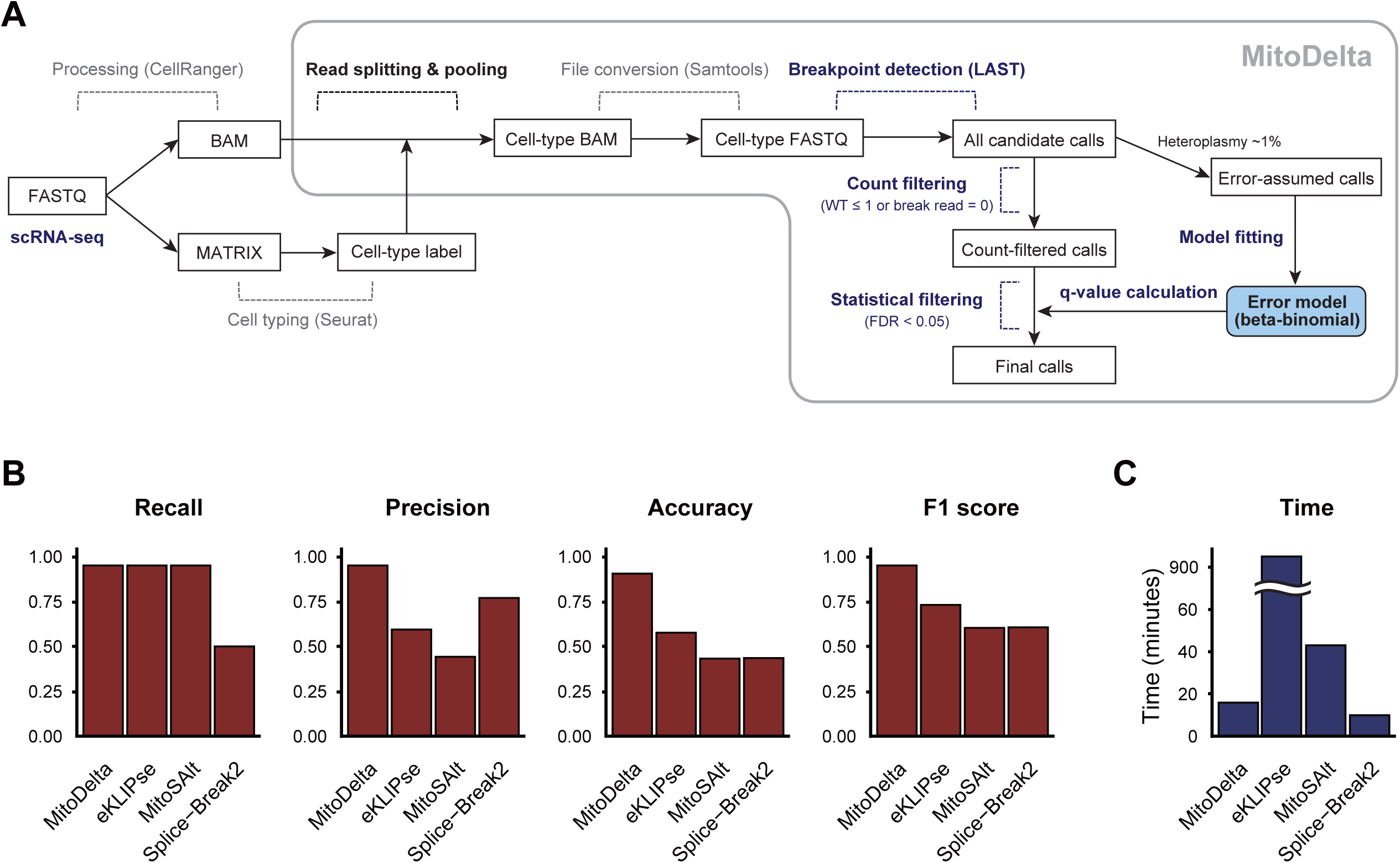
Overview of MitoDelta and benchmarking results for simulated RNA-seq data. **A** Schematic summary of the MitoDelta workflow. **B** Performance evaluation of MitoDelta, eKLIPse, MitoSAlt, and Splice-Break2 on simulated RNA-seq data. Twenty distinct mtDNA deletion variants were simulated using 50,000 mtDNA-derived reads per sample, approximating the coverage of pooled scRNA-seq data from 50 cells (assuming ∼5% mitochondrial reads and 20,000 reads per cell). Variant calling performance was assessed in terms of recall, precision, accuracy, and F1 score. **C** Total execution time for processing all 100 simulated FASTQ files (20 deletions × 5 coverage levels). MitoDelta completed the analysis in 15 minutes 55 seconds, MitoSAlt in 43 minutes 1 second, eKLIPse in 15 hours 44 minutes 21 seconds, and Splice-Break2 in 9 minutes 58 seconds.

### Benchmarking MitoDelta and existing tools on simulated scRNA-seq data

MitoDelta was compared with MitoSAlt, eKLIPse, and Splice-Break2. Although they were originally developed for genomic DNA-seq, we included them as a comparison because no existing mtDNA deletion callers have been designed for scRNA-seq data. Each method employs a different algorithmic approach and has strengths. MitoSAlt first reduce input read by premapping, and then detects deletions using split reads aligned by LAST [27]. This tool is characterized by its high sensitivity and precise breakpoint resolution. eKLIPse identifies breakpoints through a BLASTN [38] local alignment of soft-clipped reads initially mapped with BWA-MEM. This tool is designed with a focus on sensitivity and reports the sequences flanking the breakpoints, which is motivated by the biological observation that mtDNA deletions often occur between short, repeated nucleotide motifs. Splice-Break2 relies on MapSplice [39], an RNA mapper, and leverages the resulting junction reads as evidence for deletions By comparing these tools, we aimed to clarify their strengths and limitations.

For the simulated dataset, we benchmarked MitoDelta against existing tools. At our target coverage of 50,000 reads, representing 50-cell pooling, MitoDelta achieved the highest precision (0.95) and F1 score (0.95) (Fig. 1B). MitoDelta, MitoSAlt, and eKLIPse all achieved strong recall (0.95), successfully identifying most true deletions. Splice-Break2 showed lower recall, but it exhibited the second-highest precision after MitoDelta. Across all coverage levels, MitoDelta consistently maintained the highest precision (Supplementary Fig. S1). Although the recall decreased at ultralow read depths (5,000 and 10,000 reads), this trade-off was acceptable, as MitoDelta is designed for a cell-type pooled analysis, which typically provides greater coverage. Overall, the read depth had a clear negative impact on performance.

In terms of the execution time, Splice-Break2 was the most time-efficient, completing the analysis in 9 minutes 58 seconds (Fig. 1C), followed by MitoDelta (15 minutes 55 seconds) and MitoSAlt (43 minutes 1 second). In contrast, eKLIPse showed the longest runtime (15 hours 44 minutes 21 seconds). This is likely attributable to its internal function of searching for repeat sequences at the breakpoint [23].

In summary, MitoDelta outperformed the other methods on the simulated RNA-seq dataset with 50,000 or more reads, showing balanced performance between precision and recall.

### Qualitative and quantitative evaluations of MitoDelta in detecting deletions in simulated RNA-seq data

We next conducted more detailed benchmarking on the same simulated RNA-seq dataset, focusing on both qualitative and quantitative aspects. We identified critical steps for accurate deletion detection and explored areas for potential improvement by systematically modifying individual components of each tool’s workflow and assessed their impacts (Fig. 2A).

**Figure 2.**
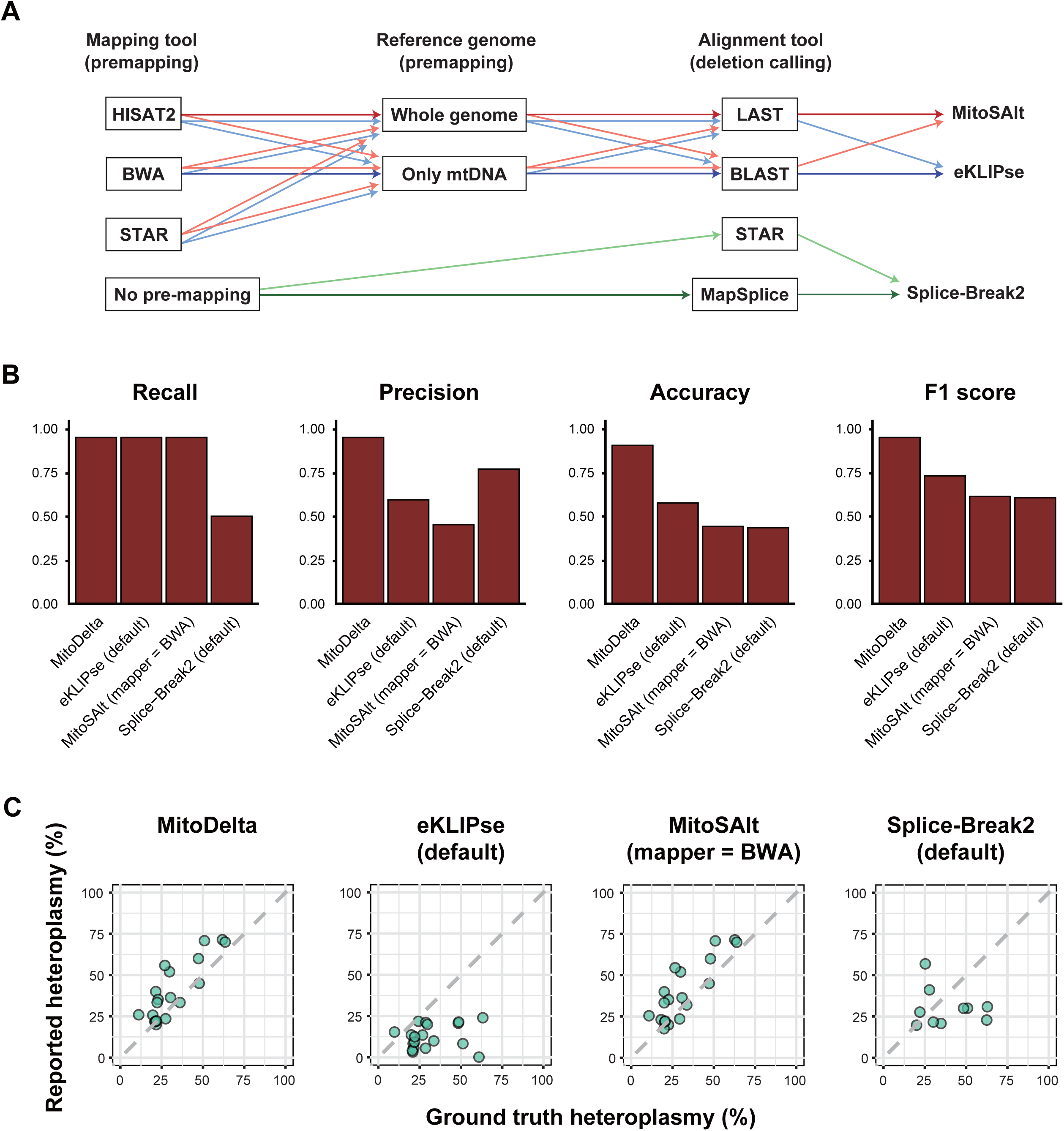
Performance evaluation of mtDNA variant callers on simulated RNA-seq data. **A** Schematic overview of the customized workflows evaluated for each variant caller. The diagram illustrates which tools were substituted at specific steps to create and test alternative workflows. The thick arrows indicate the default settings, whereas the lighter arrows represent custom workflow versions. For eKLIPse and MitoSAlt, 12 workflow combinations were generated by testing three mapping tools for premapping (BWA-MEM2, HISAT2, and STAR), two types of premapping references (mtDNA-only or whole genome), and two local alignment tools (BLAST or LAST). For Splice-Break2, two mapping strategies (MapSplice and STAR) were tested, resulting in two workflow versions. **B** Comparison of the deletion detection performance of MitoDelta, eKLIPse, MitoSAlt, and Splice-Break2. Bar plots show the recall, precision, accuracy, and F1 score for the best-performing workflow of each tool on the 50,000-read simulated dataset. The results from other coverage levels are provided in Supplementary Fig. S2. **C** Heteroplasmy estimates from the best-performing workflow (based on the F1 score) for each tool using the 50,000-read dataset. Scatter plots compare the estimated versus true heteroplasmy levels. The x-axis represents the ground truth heteroplasmy, whereas the y-axis indicates the heteroplasmy levels reported by each method. Perfect concordance is represented by the diagonal line. The full results across additional coverage levels are shown in Supplementary Figs. S3–6.

In the qualitative evaluation, MitoDelta outperformed all the other methods (Fig. 2B, Supplementary Fig. S2). While eKLIPse and Splice-Break2 achieved their best F1 scores with the default settings, MitoSAlt improved when its default mapper (HISAT2) was replaced with BWA. eKLIPse failed to detect any deletions when a premapping step was performed with HISAT2 or STAR, likely due to the lack of soft-clipped reads. Splice-Break2 showed improved recall when STAR was used instead of MapSplice, but this gain was accompanied by a decrease in precision.

For the quantitative evaluation, we assessed the accuracy of the heteroplasmy estimation. Both MitoDelta and MitoSAlt reported estimates closely aligned with the true heteroplasmy levels, whereas eKLIPse consistently underestimated them (Fig. 2C, Supplementary Figs. S3–5). Compared with MitoSAlt, MitoDelta effectively filtered out calls whose estimated heteroplasmy levels deviated from the ground truth (Supplementary Figs. S3, S5). In contrast, Splice-Break2 showed inconsistent trends, with both over- and underestimation observed (Supplementary Fig. S6). A possible explanation is its reliance on overall benchmark coverage as the background depth, an assumption that does not account for the inherently uneven coverage typical of RNA-seq data, unlike the uniform coverage of WGS or DNA-seq data.

In summary, MitoDelta had the best overall performance, achieving the highest F1 score and the most accurate heteroplasmy estimates for the simulated datasets.

### Qualitative and quantitative evaluations of MitoDelta in detecting deletions in real scRNA-seq data

We also evaluated the tools’ performance on real scRNA-seq data. The benchmark dataset included samples of PBMCs from three patients, each carrying a distinct mtDNA deletion variant. The ground truth heteroplasmy levels for these patients, as reported in the original study, were 41% (Patient 1), 66% (Patient 2), and 54% (Patient 3). We also confirmed the presence of deletions at the transcript level by visualizing the scRNA-seq data in a genome browser, and the transcript-level heteroplasmy was manually curated to be 70%, 16%, and 40% for Patients 1, 2, and 3, respectively (Supplementary Fig. S7). Notably, these observations revealed a discrepancy between the transcript-level deletion frequencies and the mtDNA heteroplasmy levels reported in the original study. Therefore, in this study, we used both transcript-level and mtDNA-level heteroplasmy as reference values for the performance evaluation.

For the real dataset, MitoDelta showed the best overall performance, with the highest F1 score of 0.75 at the representative coverage level of 1,000,000 reads, corresponding to a pooled input of 50 cells (Fig. 3A, Supplementary Fig. S8). Although the recall of MitoDelta was equal to or slightly lower than that of MitoSAlt, its high precision and accuracy resulted in the highest F1 score. At lower coverage levels (20,000–200,000 reads), corresponding to single-cell to 10-cell pooling, the recall performance of MitoDelta decreased. However, MitoDelta consistently exhibited robust performance at coverage levels corresponding to pools of 50 cells or more, which are considered practical for its application.

**Figure 3.**
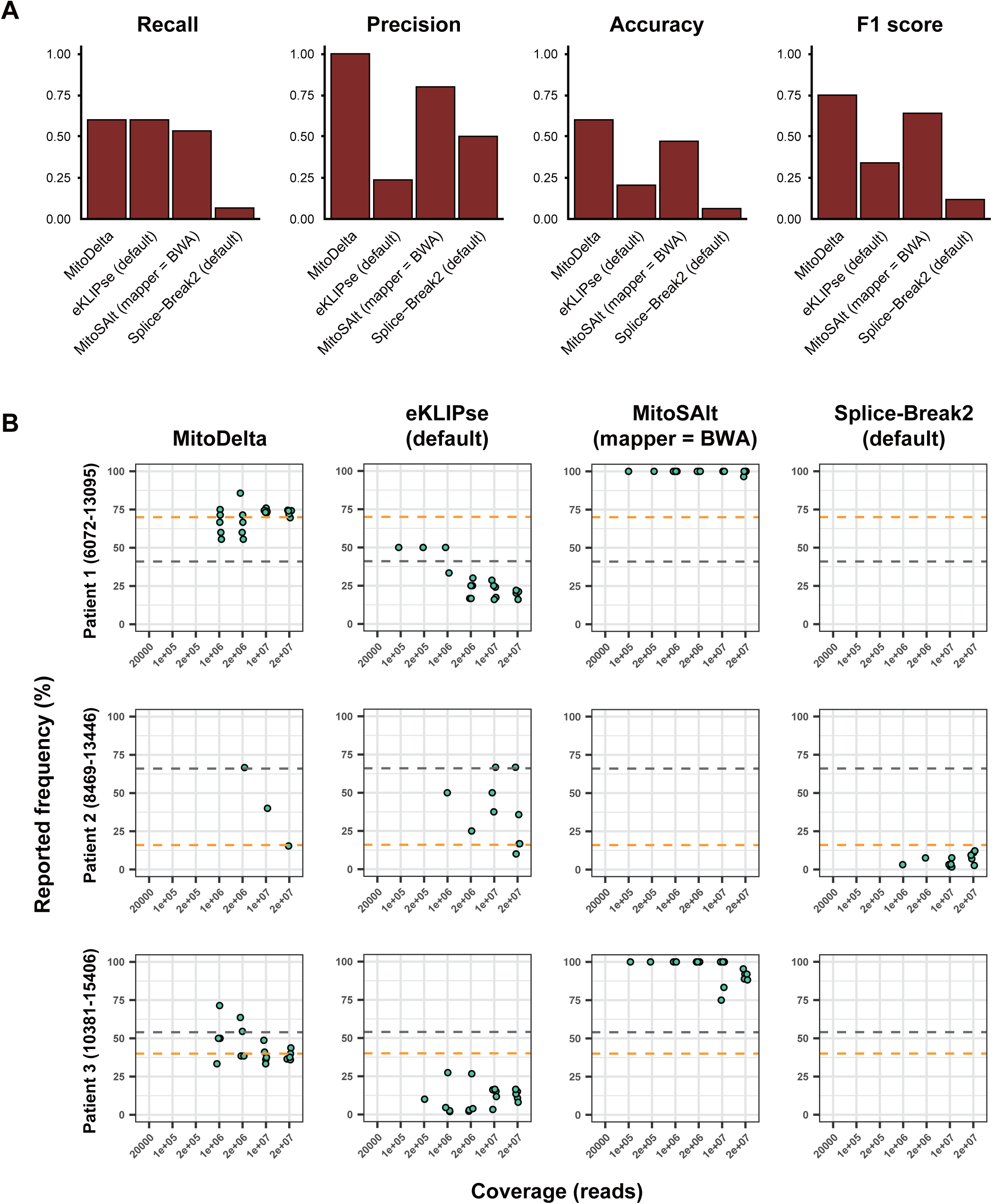
Performance evaluation on a real scRNA-seq dataset from Pearson syndrome patients. **A** Deletion detection performance for MitoDelta, eKLIPse, MitoSAlt, and Splice-Break2. Bar plots show the recall, precision, accuracy, and F1 score for the best-performing workflow of each tool evaluated using real scRNA-seq data (1,000,000 reads per sample). For eKLIPse and Splice-Break2, the default workflows showed the best performance. For MitoSAlt, a customized version using BWA as the mapping tool outperformed the default version. The results from other coverage levels and workflows are shown in Supplementary Fig. S8. **B** Heteroplasmy estimates across input coverage levels. Scatter plots show the heteroplasmy levels reported by the best-performing workflow (based on the F1 score) for each tool across various input read coverages (20,000 to 20,000,000 reads). Each coverage level includes five replicates generated by random downsampling. The x-axis represents the input read coverage, and the y-axis represents the reported heteroplasmy levels. The gray dashed line represents the heteroplasmy levels observed in the real scRNA-seq data, whereas the orange dashed line represents the mtDNA heteroplasmy values reported in the original study. The full results across all workflows are provided in Supplementary Figs. S9–11.

In terms of the heteroplasmy estimation accuracy, both MitoDelta and MitoSAlt outperformed the other tools when compared against the heteroplasmy levels inferred from scRNA-seq data, although their estimates deviated from the ground truth mtDNA heteroplasmy values determined by DNA-based analysis. MitoDelta was also effective at suppressing variant calls that deviated from scRNA-seq-level heteroplasmy (Fig. 3B, Supplementary Figs. S9–11). Consistent with the simulation results, eKLIPse tended to underestimate heteroplasmy levels, whereas Splice-Break2 failed to detect certain variants entirely. In summary, an analysis of real scRNA-seq data revealed a key limitation in the heteroplasmy estimation: while MitoDelta can provide reasonably accurate estimates of heteroplasmy reflected at the transcript level, these values are not necessarily equal to the heteroplasmy levels in mtDNA. This discrepancy is caused by a mismatch between the abundance of deletion-bearing transcripts and the underlying mtDNA deletion frequencies.

Collectively, benchmarking with simulated and real datasets revealed that MitoDelta is well suited for an mtDNA deletion analysis using scRNA-seq data, offering balanced precision and recall. It demonstrated the most robust performance in pooled scRNA-seq settings, while detection at the single-cell level appeared to be challenging due to its inherently sparse nature, even within the highly transcribed mtDNA region. An accurate estimation of mtDNA heteroplasmy was also limited by discrepancies between transcript-level deletion frequencies and actual mtDNA heteroplasmy levels. In summary, MitoDelta provides an accurate approach for estimating RNA-reflected mtDNA deletion frequencies, particularly when reads are pooled by cell type.

### Revealing cell-type-specific mtDNA deletion landscapes in Parkinson’s disease by MitoDelta

We applied MitoDelta to an snRNA-seq dataset from human postmortem brain samples of 15 patients with sporadic PD and 14 age-matched controls to demonstrate its practical value. The analysis included the following cell typing and MitoDelta steps: read pooling, variant calling, and statistical filtering (Fig. 4A).

**Figure 4.**
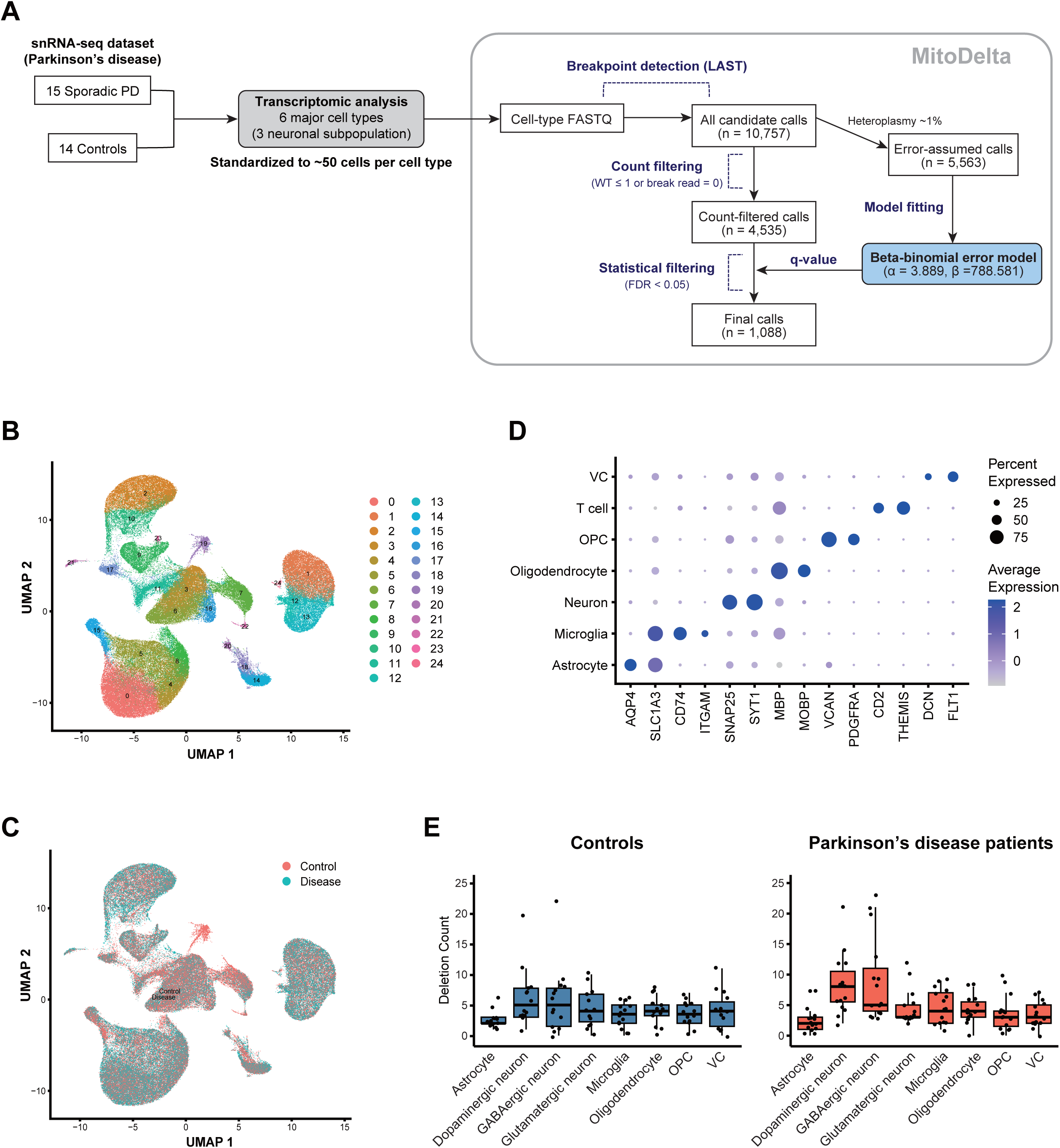
Application of MitoDelta to a snRNA-seq dataset from Parkinson’s disease patients. **A** Overview of the analysis workflow. **B** UMAP plot of 15 Parkinson’s disease (PD) and 14 control samples, revealing 25 transcriptionally distinct clusters. **C** Distribution of nuclei from PD and control samples across the clusters. **D** Expression patterns of representative marker genes used to annotate major cell types. **E** Cell-type-specific mtDNA deletion variant counts estimated by MitoDelta. This analysis was performed on downsampled data (a maximum of 50 nuclei per cell type per individual). No significant differences in the deletion burden were observed across cell types in the control samples (KruskalLWallis test, p = 0.17). In PD samples, significant variation was observed across cell types (KruskalLWallis test, p < 0.001), with dopaminergic and GABAergic neurons showing elevated deletion counts (Dunn’s test, p < 0.05).

In the cell typing process, unsupervised clustering identified 25 distinct clusters (Fig. 4B). The cells from both the PD and control samples were broadly distributed across the clusters, except for Cluster 19 (Fig. 4C). Based on marker gene expression, these clusters were classified into seven major cell types: astrocytes, microglia, neurons, oligodendrocytes, OPCs, T cells, and VCs (Fig. 4D; Supplementary Fig. S12). Neurons were further subclassified into dopaminergic, GABAergic, and glutamatergic subtypes based on differentially expressed gene signatures. The cell population sizes varied by cell type (Supplementary Fig. S13), with the distributions largely consistent with those of the original study, supporting the validity of our cellular annotations. Cell populations that were too small, such as Cluster 19 and T cells, were excluded from subsequent analyses. The number of cells per cell type was standardized by downsampling to a maximum of 50 cells to ensure comparability across samples and cell types.

The resulting pooled reads were then processed with MitoDelta (Fig. 4A). The initial variant calling step detected 10,757 candidate mtDNA deletions. Of these, 5,563 ultralow heteroplasmy variants (∼1%) were assumed to originate from background sequencing errors and were used to fit a beta-binomial error model. Using the fitted model, we calculated the p value of each variant as being error derived. Next, we excluded variants supported by only a single breakpoint read or lacking wild-type reads, leaving 4,535 candidates for the following evaluation. Multiple testing correction was performed using the BenjaminiLHochberg procedure, and q values were calculated. Applying a false discovery rate (FDR) threshold of 0.05, we retained 1,088 high-confidence deletion calls as statistically significant (Supplementary Table S1).

Using this result, we validated the performance of MitoDelta from two perspectives: (1) its ability to identify common mtDNA deletions and (2) its ability to resolve cell-type-specific deletion patterns. First, we focused on a well-known common deletion variant in PD patients (chrM: 8469-13447) [40]. MitoDelta successfully identified this deletion in dopaminergic neurons from one PD patient (s.0118) (Supplementary Table S1), and its presence was also visually confirmed using a genome browser (Supplementary Fig. S14). Next, we explored mtDNA deletion patterns at a cell-type resolution. In PD samples, deletion burdens varied significantly across cell types (KruskalLWallis test, p < 0.001), with dopaminergic and GABAergic neurons exhibiting greater deletion burdens than other cell types (Dunn test, p < 0.05) (Fig. 4E). In contrast, no significant differences were observed in the control samples (KruskalLWallis test, p = 0.17). These findings not only align with previous reports of mtDNA deletions in dopaminergic neurons in PD patients [14] but also indicate an increased burden in GABAergic neurons, which has not been widely reported.

## Discussion

In this study, we present MitoDelta, a computational pipeline for identifying mtDNA deletion variants at cell-type resolution from scRNA-seq data. Although several SNV callers exist for scRNA-seq data [20], mtDNA deletions have been largely overlooked in this context. MitoDelta achieved this goal by combining a transcriptional analysis, sensitive alignment, and statistical refinement. This integrated approach allows for balanced recall and precision, enabling a reanalysis of scRNA-seq data without requiring additional experimental procedures.

Most existing mtDNA deletion callers are designed for DNA-seq data and thus have limited applicability to scRNA-seq data. However, mtDNA-derived reads are relatively dense in scRNA-seq data compared with nuclear transcripts due to the high copy number, stable expression, and coding nature of mitochondrial transcripts. By leveraging these reads as “byproducts,” we found that deletion variants, particularly those within the mitochondrial genome, can be detected even from scRNA-seq data using sensitive alignment strategies.

MitoDelta has three key strengths. First, it enables the detection of mtDNA deletion variants at cell-type resolution. Unlike conventional DNA-based methods, which typically lack a transcriptomic context, MitoDelta allows for the simultaneous capture of transcriptional profiles and mtDNA deletions. This approach leads to a more integrated understanding of cellular states and mitochondrial variant dynamics. Second, MitoDelta effectively reduces false-positive calls by incorporating a beta-binomial model during its filtering step. This statistical approach is well suited for modeling overdispersed data, such as RNA-seq or low-abundance DNA-seq data [30,31], thereby leading to more reliable variant calls. Third, MitoDelta is compatible with a wide range of CB-based scRNA-seq platforms, such as 10x Genomics Chromium, Drop-seq [41], and Quarz-seq2 [42].

We reanalyzed a published single-nucleus transcriptomic dataset from patients with Parkinson’s disease to demonstrate the usefulness of MitoDelta. Our analysis revealed increased mtDNA deletion burdens in neurons, consistent with previous reports [14,43,44]. In these studies, cell-type-resolved analyses were conducted using cell isolation techniques, such as cell sorting or laser-capture microdissection. These methods have provided valuable insights into specific cell types but are generally limited in throughput and lack transcriptomic data. In contrast, MitoDelta enables a high-throughput and transcriptome-integrated approach involving the use of single-cell RNA-seq data. For example, previous cell sorting-based studies have identified mtDNA deletions in dopaminergic neurons [14], but the deletion patterns in other subtypes have remained unclear. In our study, we reaffirmed the high deletion burden in dopaminergic neurons. Dopaminergic neuron loss in the midbrain is a hallmark of PD, and these neurons are especially vulnerable in PD patients [45,46]. Both previous studies and our analysis revealed substantial mtDNA deletions in dopaminergic neurons, suggesting that such genomic alterations may underlie their selective vulnerability in PD patients. In addition, MitoDelta identified a high deletion burden in GABAergic neurons, a cell population that was not previously reported to be associated with mtDNA deletions in the context of PD. This suggested its potential for identifying previously unrecognized patterns of mitochondrial genome alterations. Together, MitoDelta represents a powerful approach to explore mitochondrial variant dynamics and corresponding transcriptomics.

Despite its advantages, MitoDelta has several limitations. First, an accurate estimation of mtDNA heteroplasmy remains challenging because of discrepancies between actual mtDNA heteroplasmy levels and those reflected in transcript reads [35]. A possible explanation is the increased instability and degradation of variant-harboring transcripts, making the RNA-level deletion frequencies different from those in DNA. Therefore, heteroplasmy estimates based solely on RNA-seq data may not fully capture true mtDNA heteroplasmy. However, transcript-level deletions still provide valuable insights, as they reflect functional output. Another limitation is its reliance on cell-type pooled analyses rather than single-cell evaluations because the reads per single cell are too sparse for variant calling. Using techniques that provide deeper sequencing coverage per cell could address this limitation.

For our prospects, full-length RNA-seq technologies would be beneficial for improving our calling approach. For example, Smart-seq2 [47], a full-length scRNA-seq platform, provides more uniform read coverage than 10x Genomics Chromium does [20]. A well-covered read distribution could be advantageous for improving deletion detection. Similarly, other full-length approaches, such as FLASH-seq [48], RamDA-seq [49], Smart-seq3 [50], and long-read scRNA-seq (e.g., PacBio [51] or Nanopore [52]), would provide well-covered reads, which are advantageous for capturing structural variants that short reads often miss. Expanding MitoDelta to these techniques could further refine the calling for mtDNA deletion variants in single cells.

MitoDelta also holds promise in spatial transcriptomics. Since most spatial transcriptomic data share a similar format with single-cell transcriptomic data, adapting MitoDelta for spatial transcriptomics could be straightforward. Some previous studies have reported spatial differences in mtDNA deletion landscapes within tumors [53], highlighting the potential significance of mtDNA deletion analyses. MitoDelta could investigate mitochondrial genomic abnormalities in spatially resolved transcriptomic datasets, ultimately offering advantages in tumor spatial genomics.

### Conclusions

This study suggests that scRNA-seq data can be effectively used for mtDNA deletion variant calling. By integrating LAST-based calling and beta-binomial model filtering, MitoDelta successfully identifies mtDNA deletions using only scRNA-seq data. MitoDelta has been shown to be applicable to scRNA-seq data through comprehensive benchmarking and provides advantages for reanalyzing public scRNA-seq data to determine the mtDNA deletion landscape.

## Supplementary materials

Supplementary Material 1, .pdf, Figures S1 to S14

Supplementary Material 2, .xls, Table S1

## Declarations

### Ethics approval and consent to participate

Not applicable.

### Consent for publication

All the authors reviewed and consented to the publication of this manuscript.

### Availability of data and materials

The source code of MitoDelta is publicly available on GitHub (https://github.com/NikaidoLaboratory/mitodelta). The datasets supporting the conclusions of this article are provided within the article, its supplementary files, and at https://github.com/NikaidoLaboratory/mitodelta/analysis.

### Authors’ contributions

HN and IN designed the research approaches used in this study. HN conducted experiments to benchmark the tools, designed the source code for this method, and performed the analysis using this method under the supervision of IN. Y. Shima and Y. Sasagawa proposed a method for removing the nuclear-transferred mitochondrial genome from the reference genome. All the authors have read and approved the final version of the manuscript.

## Acknowledgements

We thank Yuta Kochi for providing instructions on the analytical methods. We extend our sincere appreciation to Yoshimi Iwayama, Akihiro Matsushima, and Takumi Ichikawa for their expert technical assistance. Computational resources were provided by the HOKUSAI supercomputer system at RIKEN. We would also like to express our appreciation to Masanaga Yamawaki and Eriko Okada of the Department of Medical Education Research and Development for their academic support and encouragement.

