## Supplementary Material for "MitoDelta: identifying mitochondrial DNA deletions at cell-type resolution from single-cell RNA sequencing data"

#### Figure S1

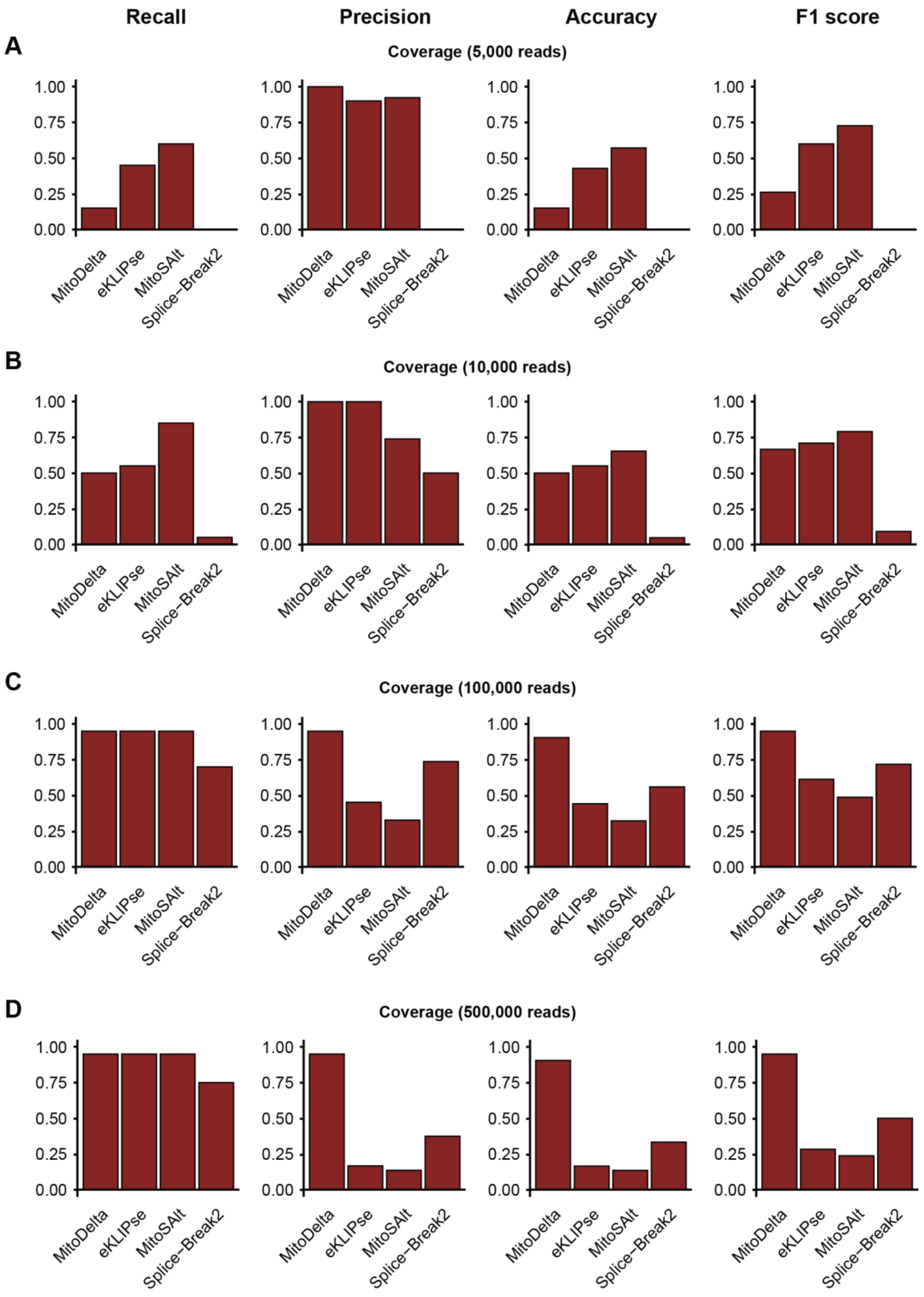


**Figure S1.** Performance evaluation of mtDNA variant callers on simulated RNA-seq data. Datasets were generated at five coverage levels, ranging from 5,000 to 500,000 mtDNA transcript-derived reads, to assess the impact of the read coverage. These levels correspond to the expected read counts from pooled samples of approximately 5 to 500 cells. The benchmarking results are shown for four representative coverage levels: **A** 5,000, **B** 10,000, **C** 100,000, and **D** 500,000 reads.

### Figure S2

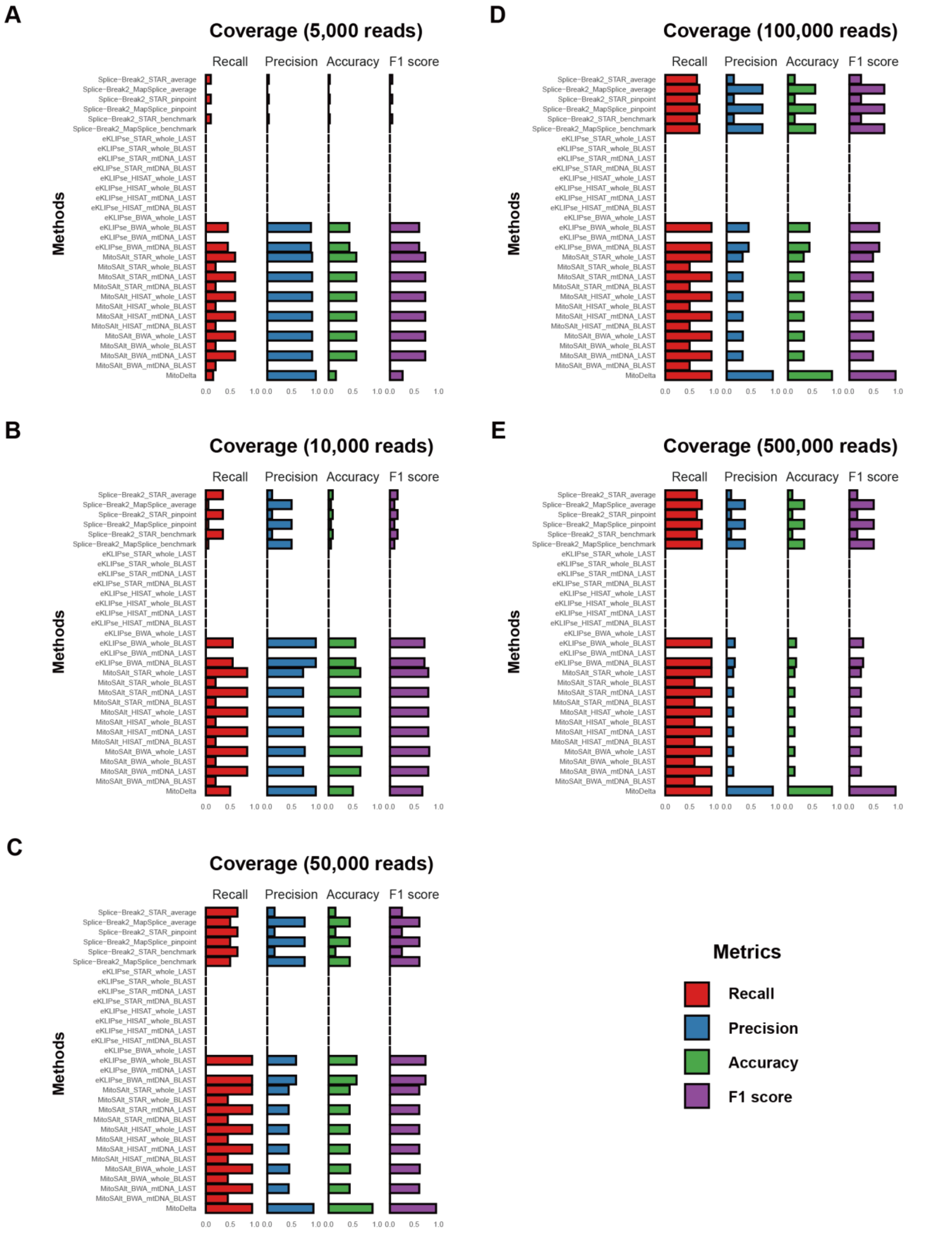


**Figure S2**. Performance evaluation of MitoDelta and existing mtDNA variant callers across custom workflows using simulated RNA-seq data.

Bar plots show the performance of all tested workflows for MitoDelta, eKLIPse, MitoSAlt, and Splice-Break2 across five read coverage levels (**A** 5,000, **B** 10,000, **C** 50,000, **D** 100,000, and **E** 500,000 reads). Each panel corresponds to a specific coverage level and displays four evaluation metrics: recall, precision, accuracy, and F1 score.

#### Figure S3

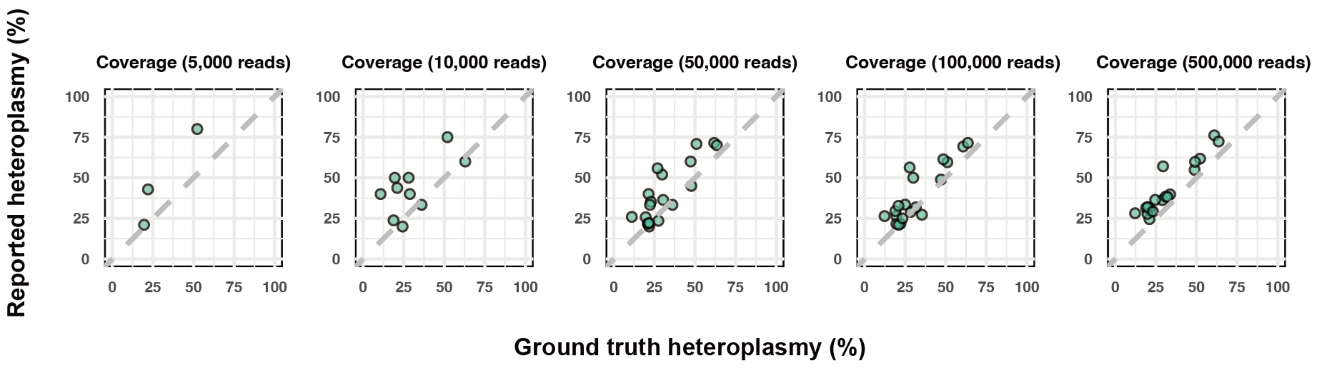


**Figure S3.** Performance of MitoDelta in estimating heteroplasmy in simulated RNA-seq data across multiple coverage levels.
Scatter plots show the concordance between heteroplasmy levels reported by MitoDelta and the ground truth values across five simulated RNA-seq datasets (5,000, 10,000, 50,000, 100,000, and 500,000 reads). The x-axis indicates the true heteroplasmy values used in the simulations, and the y-axis shows the values estimated by MitoDelta. Dots aligned along the diagonal represent perfect concordance.

### Figure S4

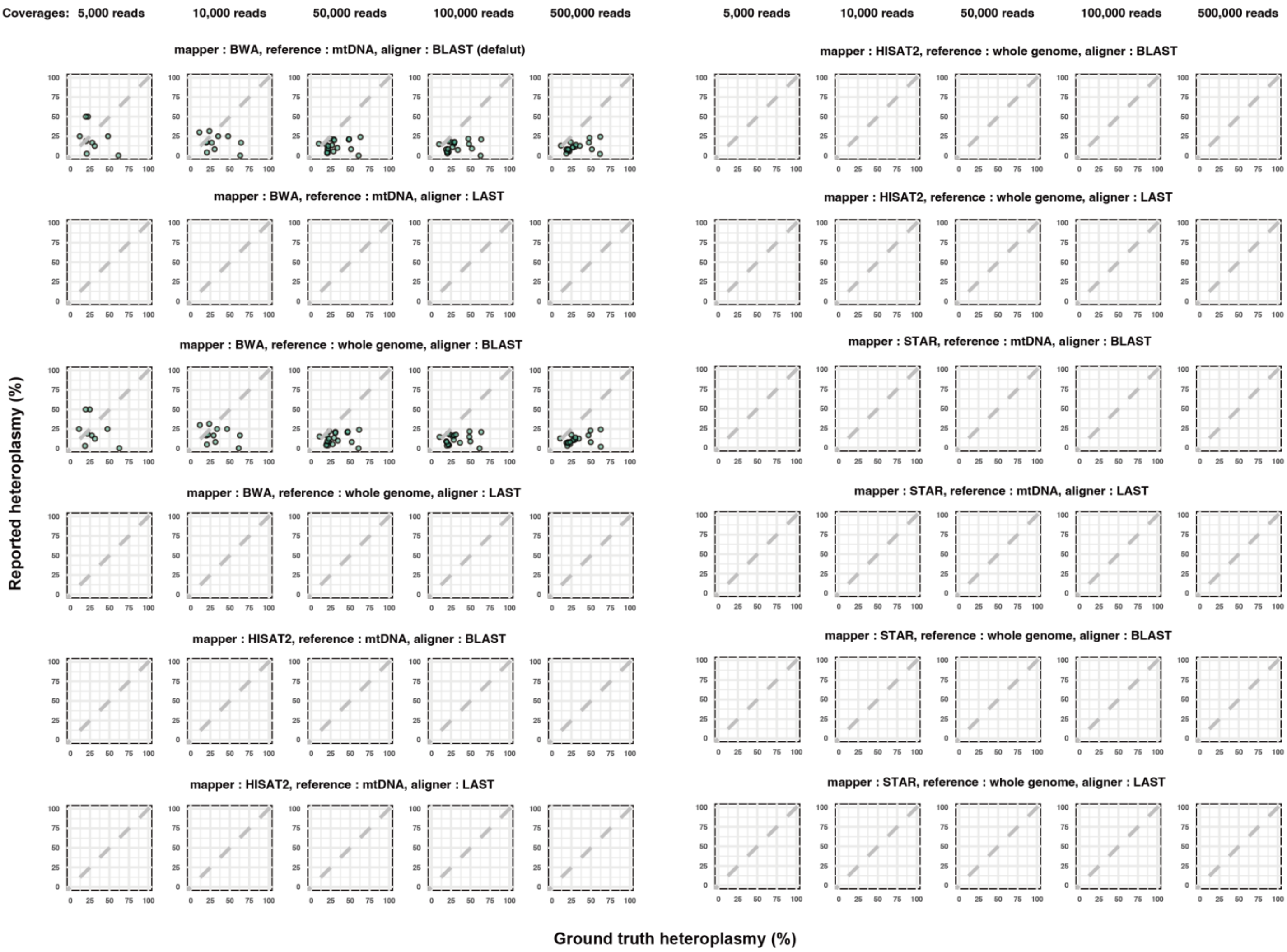


**Figure S4.** Performance of eKLIPse in estimating heteroplasmy in simulated RNA-seq data across multiple workflows and coverage levels.

Scatter plots show the concordance between heteroplasmy levels reported by eKLIPse and the ground truth values across five simulated RNA-seq datasets (5,000, 10,000, 50,000, 100,000, and 500,000 reads). The x-axis indicates the true heteroplasmy values used in the simulations, and the y-axis shows the values estimated by eKLIPse. Dots aligned along the diagonal represent perfect concordance.

### Figure S5

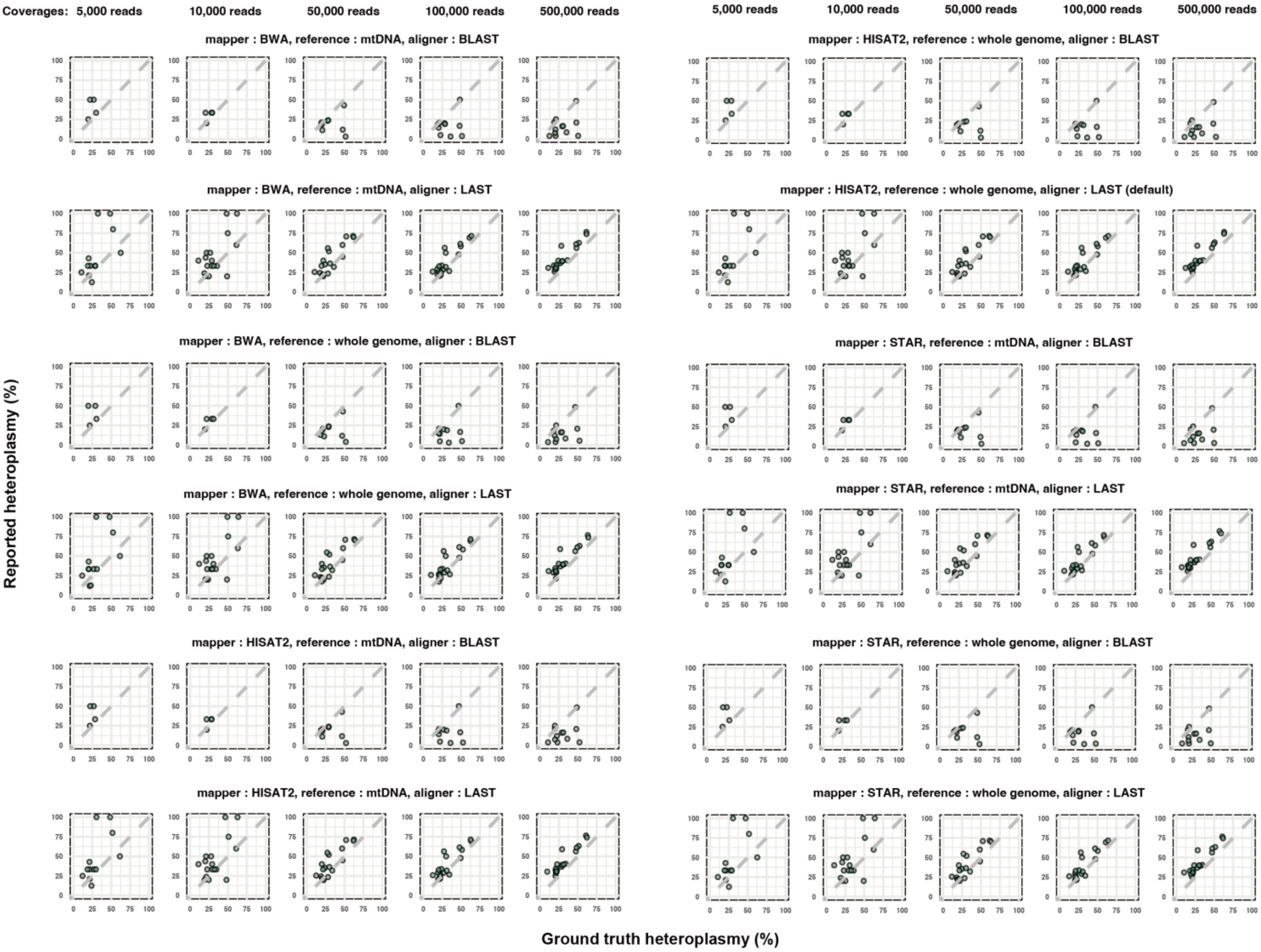


**Figure S5.** Performance of MitoSAlt in estimating heteroplasmy in simulated RNA-seq data across multiple workflows and coverage levels.

Scatter plots show the concordance between heteroplasmy levels reported by MitoSAlt and the ground truth values across five simulated RNA-seq datasets (5,000, 10,000, 50,000, 100,000, and 500,000 reads). The x-axis indicates the true heteroplasmy values used in the simulations, and the y-axis shows the values estimated by MitoSAlt. Dots aligned along the diagonal represent perfect concordance.

### Figure S6

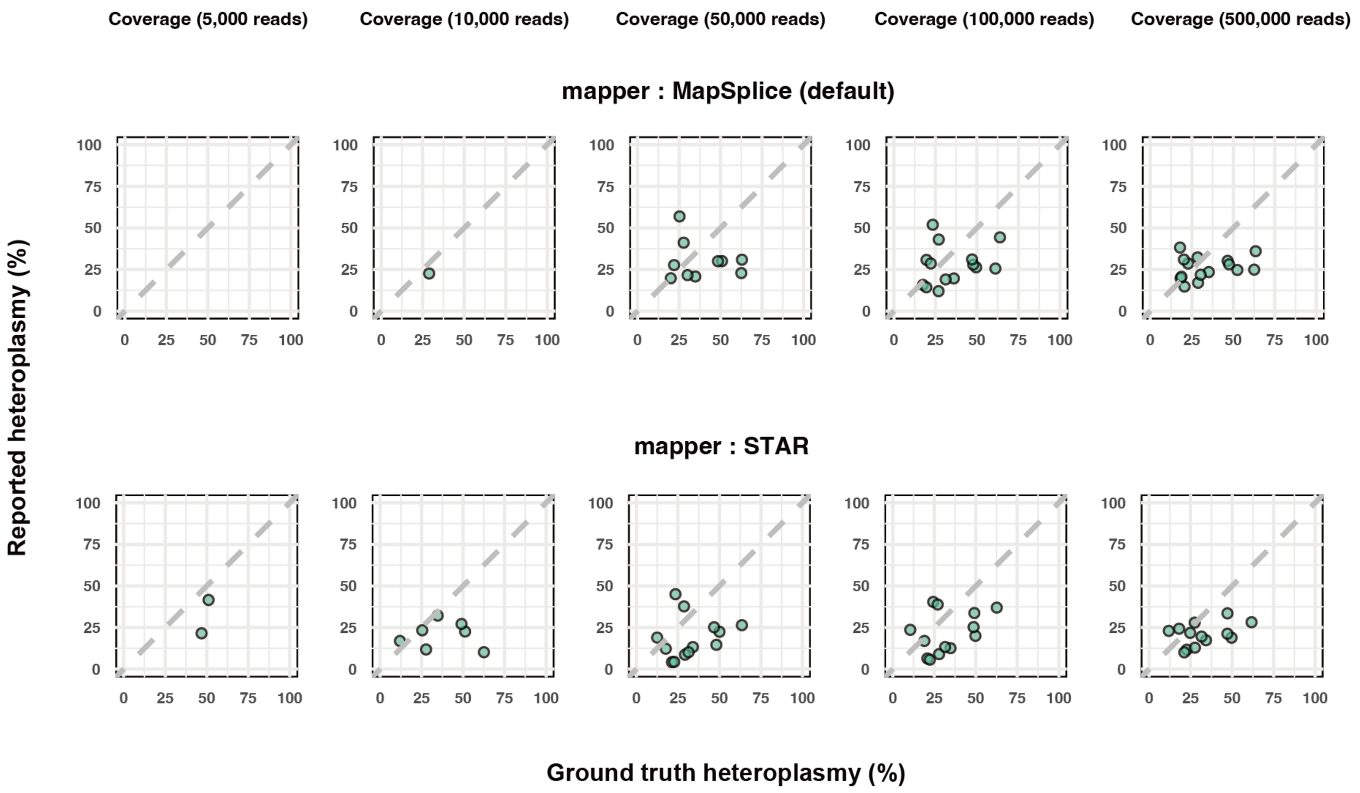


**Figure S6.** Performance of Splice-Break2 in estimating heteroplasmy in simulated RNA-seq data across multiple workflows and coverage levels.

Scatter plots show the concordance between heteroplasmy levels reported by Splice-Break2 and the ground truth values across five simulated RNA-seq datasets (5,000, 10,000, 50,000, 100,000, and 500,000 reads). The x-axis indicates the true heteroplasmy values used in the simulations, and the y-axis shows the values estimated by Splice-Break2. Dots aligned along the diagonal represent perfect concordance.

**Figure S7**

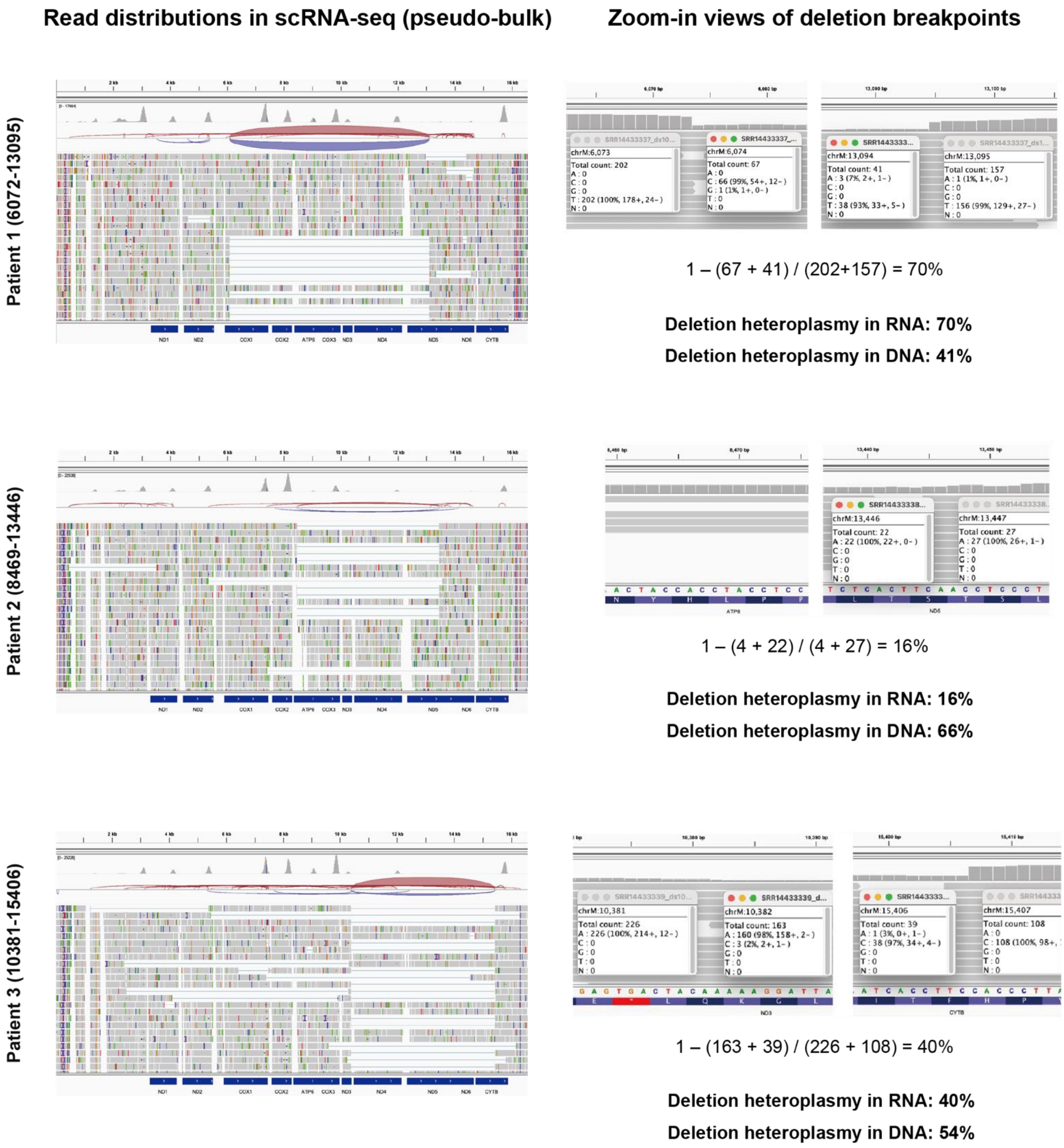


**Figure S7.** Visualization of mtDNA deletions via scRNA-seq and corresponding reference heteroplasmy levels in samples from Pearson syndrome patients.

This figure shows the scRNA-seq data used for benchmarking mtDNA deletion detection tools derived from three patients with Pearson syndrome, each harboring a unique mtDNA deletion: Patient 1 (6072–13095), Patient 2 (8469–13446), and Patient 3 (10381–15406). The left panels show overviews of the pseudobulk read distribution across the mtDNA, and the right panels display zoomed-in views of the regions surrounding each deletion breakpoint. The heteroplasmy levels in the scRNA-seq data were manually curated by calculating the ratio of the average read depth at the deletion breakpoints to that in adjacent wild-type regions. The curated levels observed in the scRNA-seq data were 70%, 16%, and 40% for Patients 1, 2, and 3, respectively. As ground truth values, we used the mtDNA heteroplasmy levels reported in the original study: 41% (Patient 1), 66% (Patient 2), and 54% (Patient 3).

### Figure S8

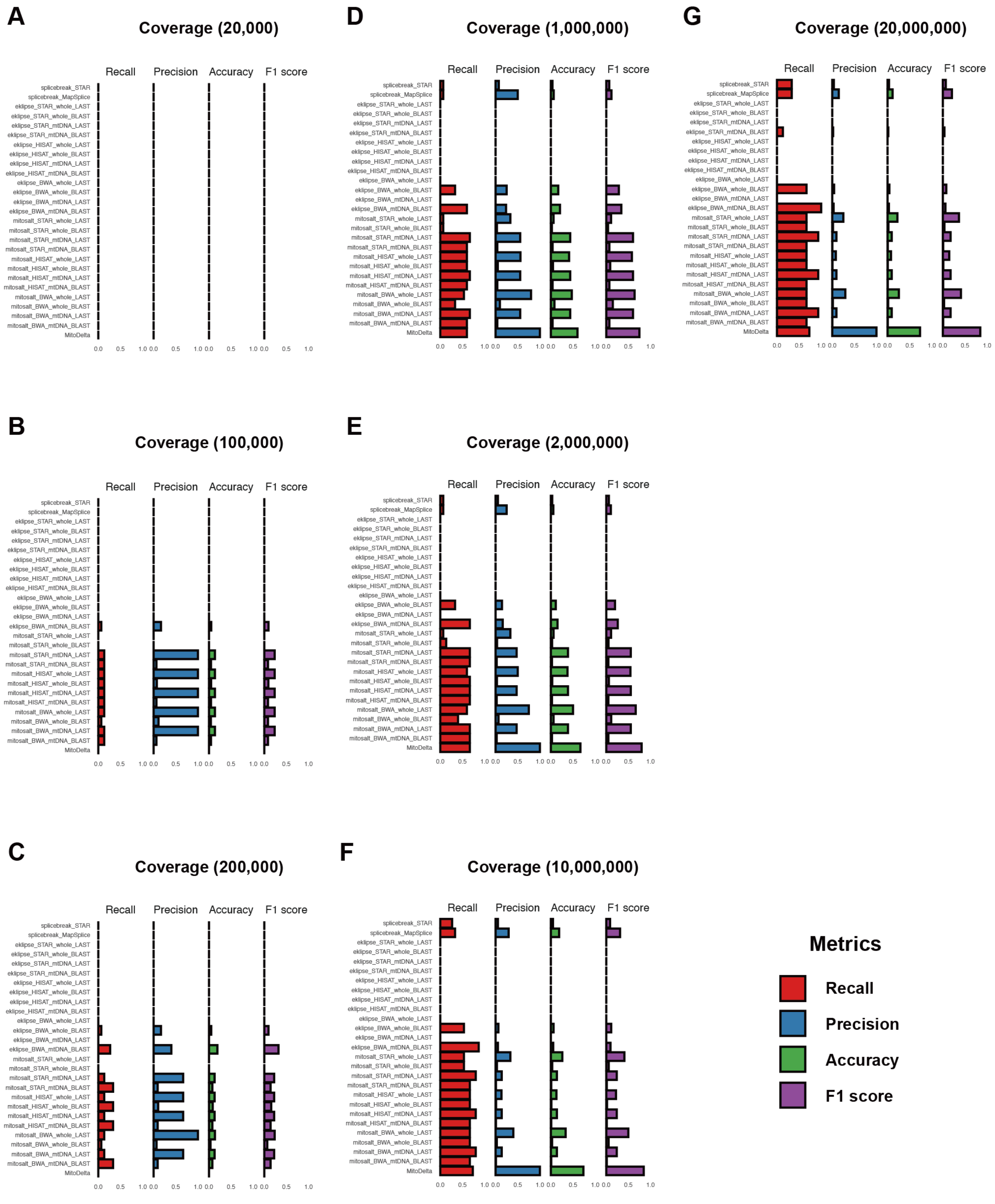


**Figure S8.** Performance evaluation of MitoDelta and existing mtDNA variant callers across custom workflows using real scRNA-seq data.
Bar plots show the performance of all tested workflows for MitoDelta, eKLIPse, MitoSAlt, and Splice-Break2 across seven read coverage levels (**A** 20,000, **B** 100,000, **C** 200,000, **D** 1,000,000, **E** 2,000,000, **F** 10,000,000, and **G** 20,000,000 reads). Each panel corresponds to a specific coverage level and displays four evaluation metrics: recall, precision, accuracy, and F1 score.

### Figure S9

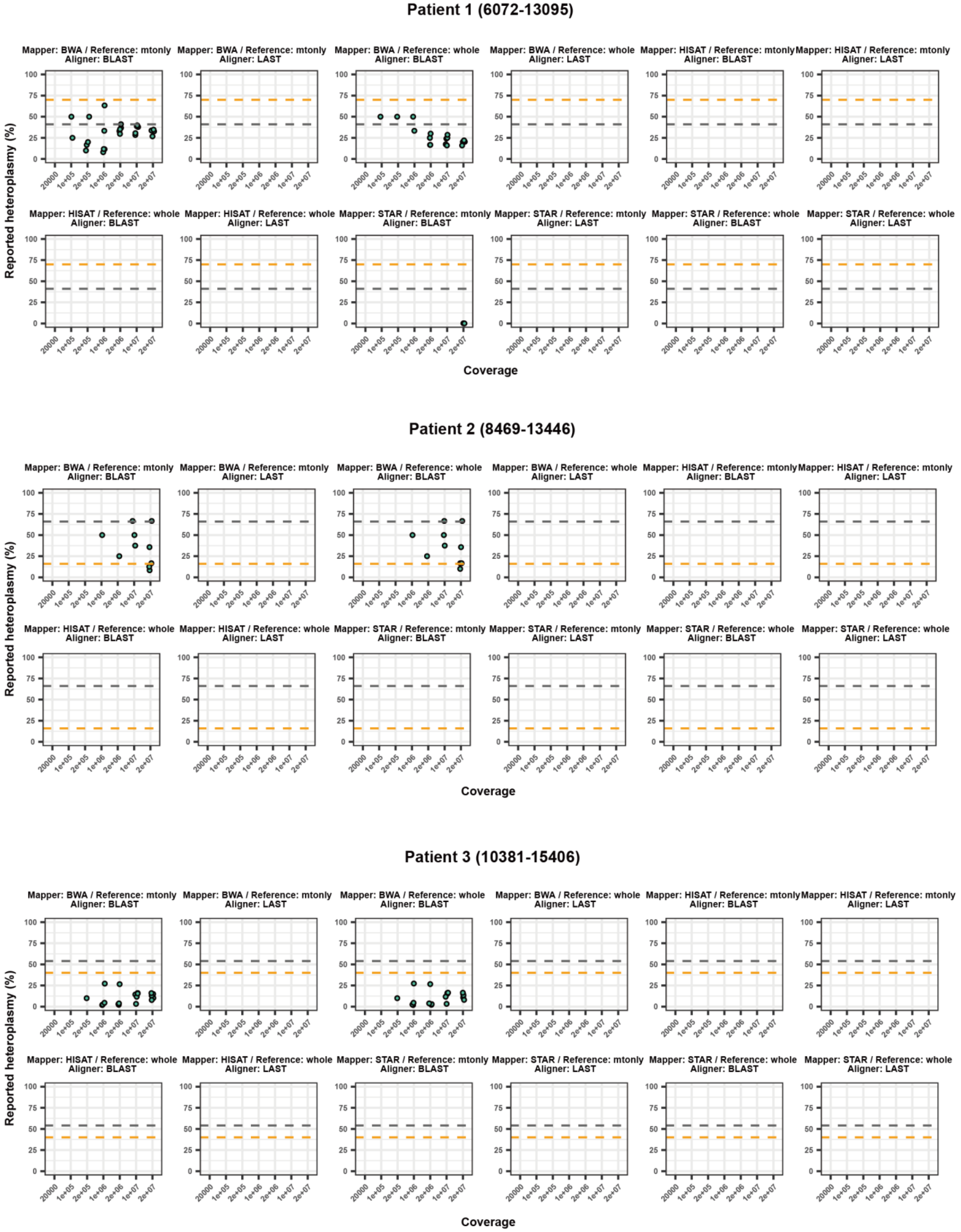


**Figure S9.** Performance of eKLIPse in estimating heteroplasmy in real RNA-seq data across multiple workflows and coverage levels.

Scatter plots compare the heteroplasmy levels reported by various eKLIPse workflows with the ground truth values in real scRNA-seq datasets. Each panel represents results from a distinct workflow. The x-axis represents the input read coverage, and the y-axis indicates the reported heteroplasmy levels. The gray dashed line represents the heteroplasmy levels observed in the scRNA-seq data; the orange dashed line represents the mtDNA heteroplasmy values reported in the original study.

### Figure S10

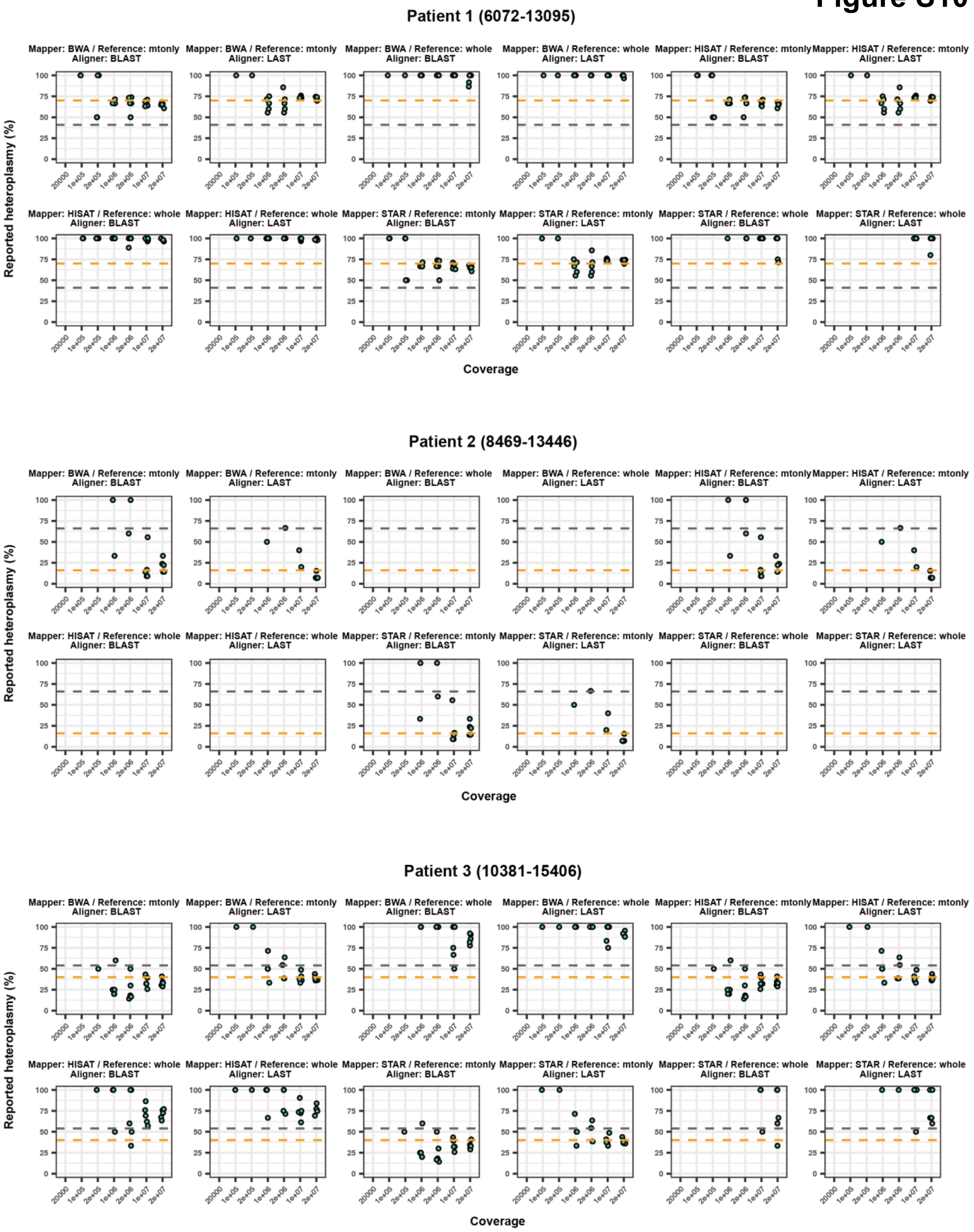


**Figure S10.** Performance of MitoSAlt in estimating heteroplasmy in real RNA-seq data across multiple workflows and coverage levels.

Scatter plots compare the heteroplasmy levels reported by various eKLIPse workflows with the ground truth values in real scRNA-seq datasets. Each panel represents results from a distinct workflow. The x-axis represents the input read coverage, and the y-axis indicates the reported heteroplasmy levels. The gray dashed line represents the heteroplasmy levels observed in the scRNA-seq data; the orange dashed line represents the mtDNA heteroplasmy values reported in the original study.

### Figure S11

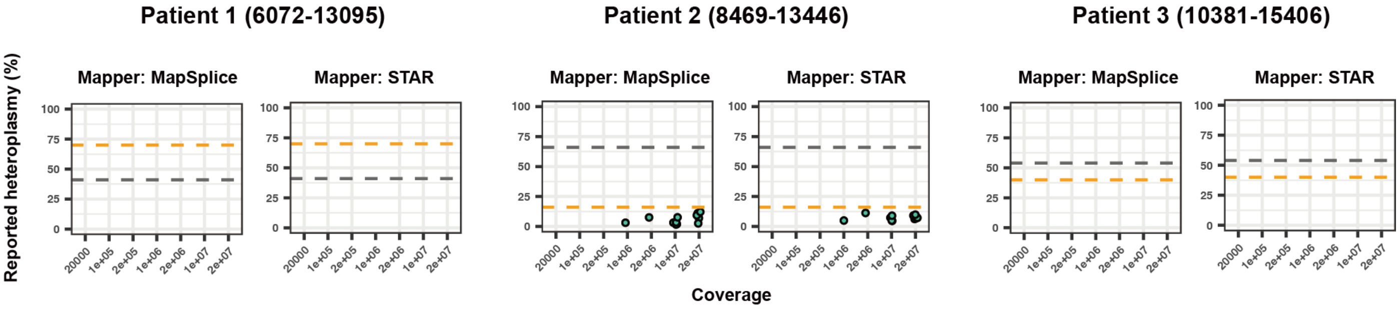


**Figure S11.** Performance of Splice-Break2 in estimating heteroplasmy in real RNA-seq data across multiple workflows and coverage levels.

Scatter plots compare the heteroplasmy levels reported by various eKLIPse workflows with the ground truth values in real scRNA-seq datasets. Each panel represents results from a distinct workflow. The x-axis represents the input read coverage, and the y-axis indicates the reported heteroplasmy levels. The gray dashed line represents the heteroplasmy levels observed in the scRNA-seq data; the orange dashed line represents the mtDNA heteroplasmy values reported in the original study.

### Figure S12

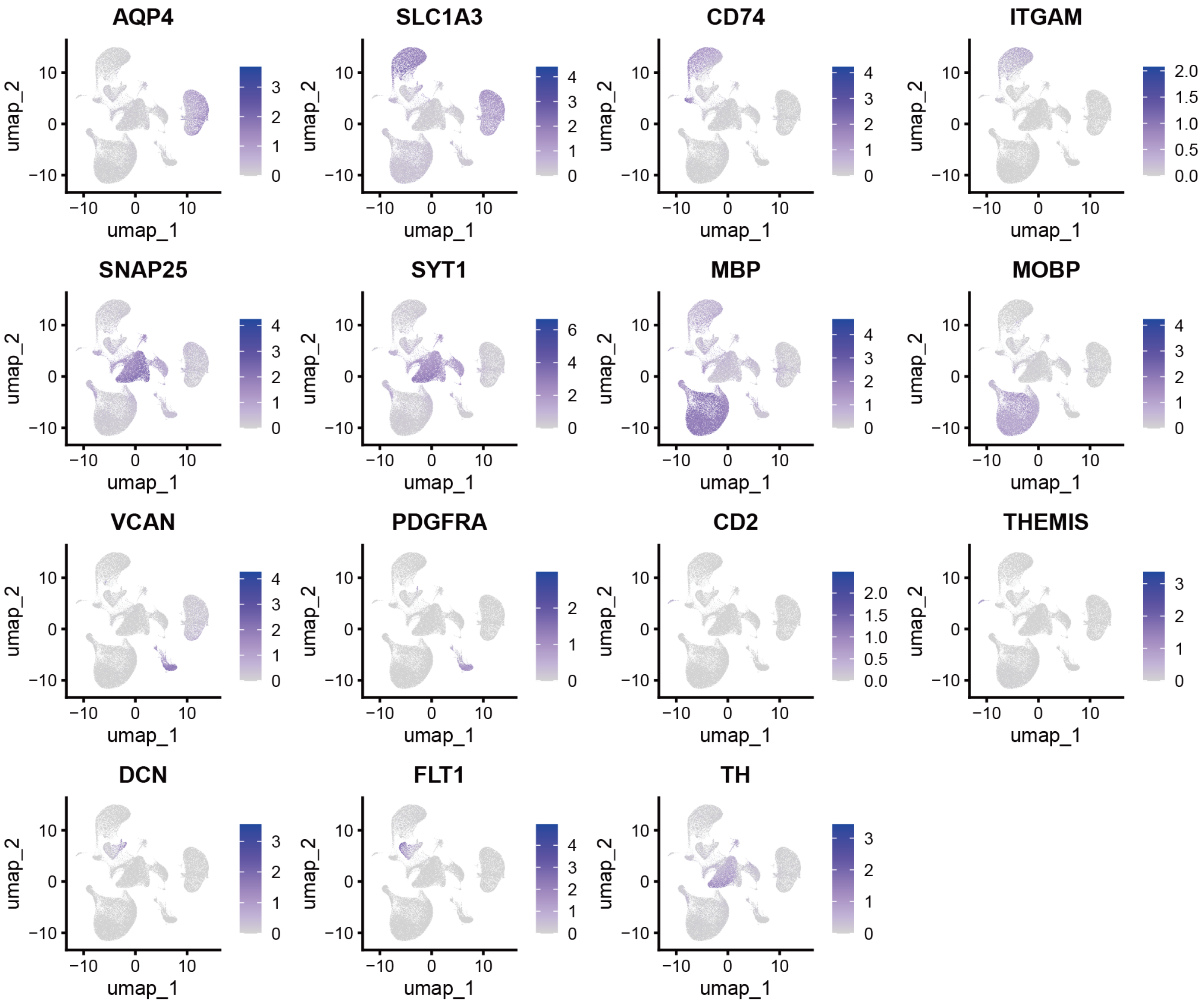


**Figure S12.**
Marker gene expression data used for the cell type annotation.
Key marker gene expression data are shown in the UMAP plot. SCT-normalized expression levels are shown instead of nFeatures.

### Figure S13

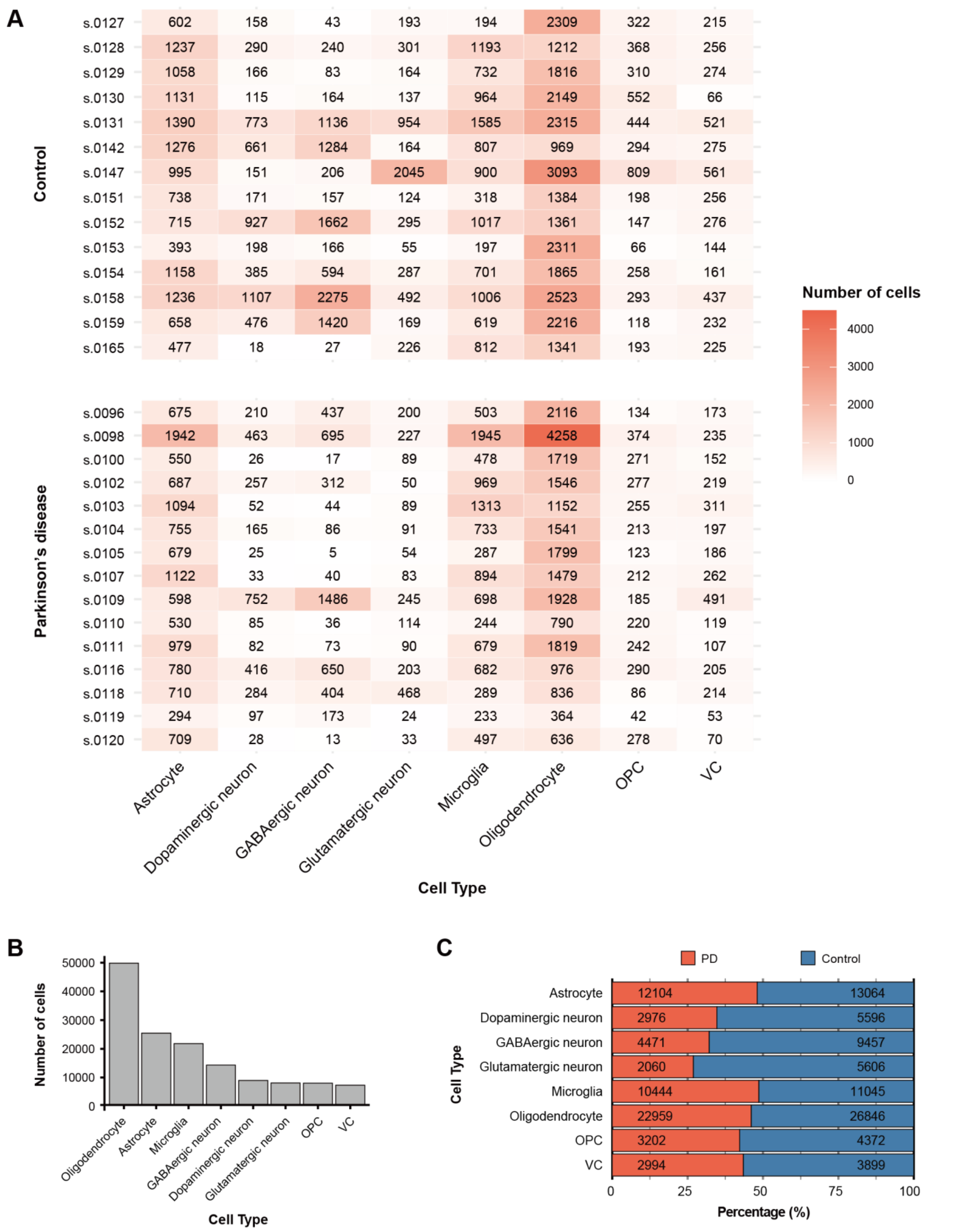


**Figure S13.** Summary of cell type distributions across samples.

**A** Table showing the number of cells for each cell type in individual patient samples.

**B** Bar plot illustrating the total number of cells for each cell type across the entire dataset.

**C** Bar plot showing the proportion of cells from the control and PD samples within each cell type.

### Figure S14

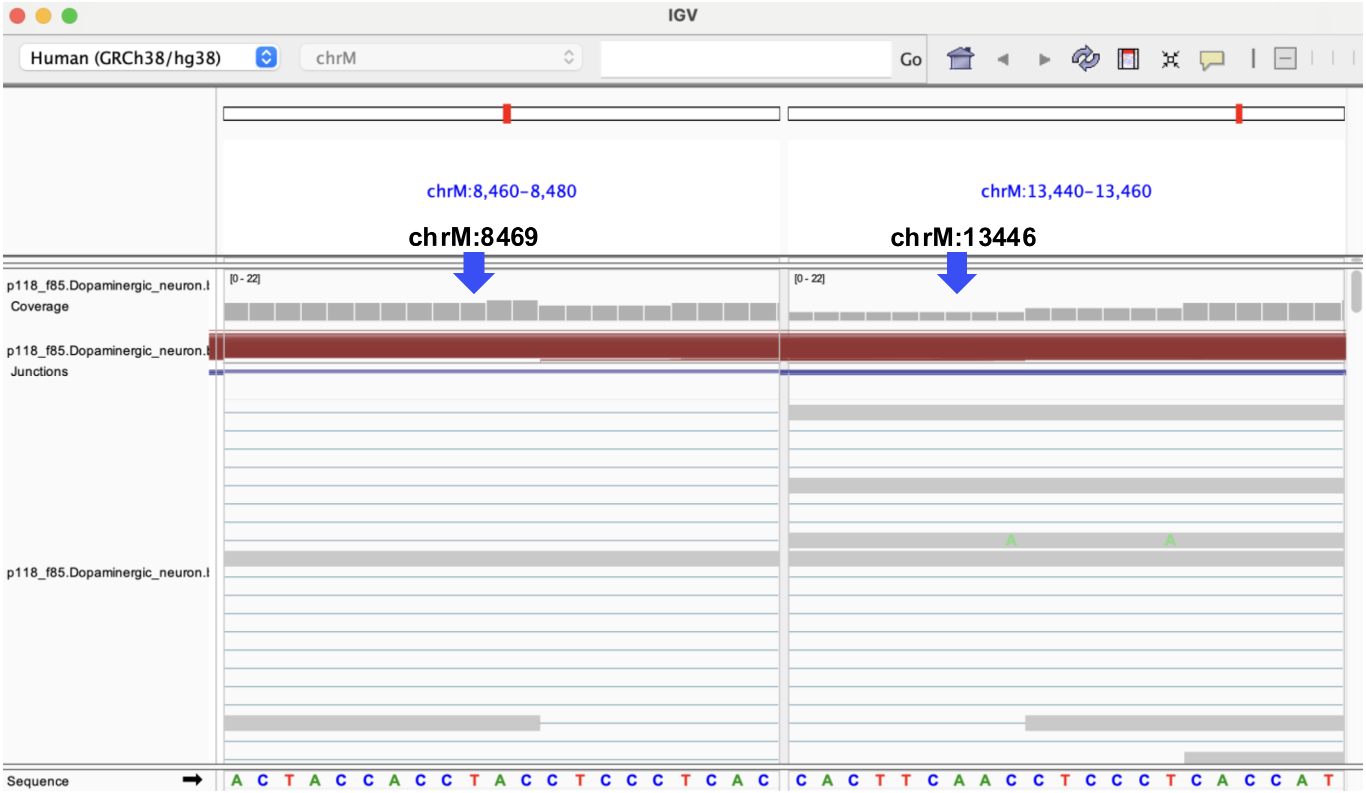


**Figure S14.** Genome browser visualization of a common mtDNA deletion identified by MitoDelta. MitoDelta identified a breakpoint (chrM: 8469–13446) in dopaminergic neurons from one PD patient (s.0118), as shown in Supplementary Table S1. This corresponds to one of the common mtDNA deletions reported in PD patients. The presence of this deletion was visually confirmed using a genome browser. Although the displayed coordinates appear to be shifted by about two bases, the surrounding sequences are identical, suggesting that the difference is due to alignment display rather than true biological variation.
